# Identification and targeting of G-quadruplex structures in *MALAT1* long non-coding RNA

**DOI:** 10.1101/2021.08.14.456336

**Authors:** Xi Mou, Shiau Wei Liew, Chun Kit Kwok

## Abstract

RNA G-quadruplexes (rG4s) have functional roles in many cellular processes in diverse organisms. While a number of rG4 examples have been reported in coding messenger RNAs (mRNA), so far only limited works have studied rG4s in non-coding RNAs (ncRNAs), especially in long non-coding RNAs (lncRNAs) that are of emerging interest and significance in biology. Herein, we report that *MALAT1* lncRNA contains conserved rG4 motifs, forming thermostable rG4 structures with parallel topology. We also show that rG4s in *MALAT1* lncRNA can interact with NONO protein with high specificity and affinity *in vitro* and in nuclear cell lysate, and we provide *in vivo* data to support that NONO protein recognizes *MALAT1* lncRNA via rG4 motifs. Notably, we demonstrate that rG4s in *MALAT1* lncRNA can be targeted by rG4-specific small molecule, peptide, and L-aptamer, leading to the dissociation of *MALAT1* rG4-NONO protein interaction. Altogether, this study uncovers new and important rG4s in *MALAT1* lncRNAs, reveals their specific interactions with NONO protein, offers multiple strategies for targeting *MALAT1* and its RNA-protein complex via its rG4 structure, and illustrates the prevalence and significance of rG4s in ncRNAs.

## INTRODUCTION

G-quadruplexes (G4s) are four-stranded nucleic acid secondary structures formed by G-rich DNA or RNA sequences, and the G-quartet layers in G4s are stabilized by monovalent ions such as potassium ion (K^+^) and sodium ion (Na^+^)(1,2). According to the classical definition, canonical G4 sequences contain four runs of guanine tracks separated by three loops (GGGN_1-7_GGGN_1-7_GGGN_1-7_GGG)(3). Recently, non-canonical G4s such as two-quartet G4s, bulged G4s and long-loop G4s were found to be prevalent in the human genome and transcriptome (4,5), highlighting the structural diversity and complexity of G4s (6,7). Studies have shown that the intervening nucleotides and the loop length can affect the stability and properties of G4 structures (8,9). Over the past decades, the structure information and the functional roles of DNA G4s (dG4s) have been studied extensively, and recent evidence has shown that they were involved in various cellular processes, including but not limited to replication, transcription, genomic instability and epigenetic regulation (10-12).

Our understanding and study of RNA G4s (rG4s) are still in its infancy (13,14). rG4s were reported to take part in numerous biological functions (14) such as microRNA (miRNAs) targeting (15,16), RNA localization (17,18), translational regulation (19,20), and alternative polyadenylation (21,22). Currently, most rG4s identified were localized in the open reading frame (ORF) (23-25) and untranslated (UTR) regions (21,26,27) of protein-coding messenger RNAs (mRNAs), which is a small fraction (< 5%) of the entire human transcriptome (28). The advent of next-generation sequencing has enabled the robust identification of non-coding RNAs (ncRNAs), which significantly facilitated our understanding and investigation of their structures and functions in cells. Interestingly, latest studies have begun to explore the existence and roles of rG4s in ncRNAs (13,14) such as long non-coding RNAs (lncRNAs) (29-33), miRNAs (34-36), Piwi-interacting RNAs (piRNAs) (37,38), ribosomal RNAs (rRNAs) (39,40), transfer RNAs (tRNAs) (41,42), and the initial findings were exciting. They revealed ncRNA rG4s to have key regulatory roles in gene expression, RNA metabolism, and protein recognition and function (13,14,43). Nevertheless, compared with mRNA G4s, which have been studied more extensively, there is still a lot of work to be done for ncRNA G4s.

RNA binding proteins (RBPs) play significant roles in regulating the RNA structure, which in turn control diverse biological processes (44,45). And rG4 structure and rG4 binding protein is no exception. Since the first rG4 binding protein, fragile X mental retardation protein (FMRP), was identified in 2001(46,47), many rG4-binding proteins have been reported to play regulatory roles by folding or unfolding the G4 structures or convening other binding motifs (25,48,49). For instance, the binding of hnRNP F to an rG4 located in the intron of CD44 promotes the inclusion of CD44 variable exon v8 and regulates alternative splicing (25). The binding of DHX36 to rG4 located at the 5’ end of the lncRNA human telomerase RNA (hTR) can unwind the RNA structure and facilitate hTR maturation and telomerase function (50,51). Moreover, the binding of nucleolin (NCL) to the rG4 in lncRNA *LUCAT1* regulates the expression of MYC thus modulate the CRC cell proliferation (49). Therefore, targeting and interfering with the rG4-protein interaction can serve as a potential strategy for manipulating gene function and regulation, and may have impact and implications in treating and preventing rG4-associated diseases.

In this study, we first identified hundreds of rG4-containing ncRNAs, and then focused on investigating rG4s on a biologically important lncRNA, *MALAT1*. We demonstrated the formation of the lncRNA *MALAT1* rG4 structures by multiple spectroscopic assays and found *MALAT1* rG4s to be highly conserved using comparative sequence analysis. Besides, we reported for the first time the specific binding between *MALAT1* rG4s and NONO protein *in vitro* and in nuclear lysate and provided substantial evidence of their *in vivo* interaction through rG4 motifs. Furthermore, to explore the targeting approach and potential application, we showed that the *MALAT1* rG4-NONO protein interaction could be targeted by rG4-specific small molecule pyridostatin (PDS), peptide RHAU53, or our recently developed rG4-targeting L-RNA aptamer (L-Apt.4-1c), leading to the dissociation of *MALAT1* rG4-NONO protein interaction.

## MATERIAL AND METHODS

### Oligonucleotide, protein, peptide, and small molecule preparation

The 5’ FAM-labeled and 5’ Biotin-labelled oligonucleotides (oligos) used in this study were synthesized by Integrated DNA Technologies (IDT). The primers used for qPCR were synthesized by Genewiz Biotechnology Co., Ltd. The L-Apt.4-1c aptamer used in this study was synthesized by ChemGenes Corporation. They were dissolved to a concentration of 100 µM (according to the supplier’s instruction) with ultra-pure nuclease-free distilled water (Thermo). All the dissolved oligos were stored at -20 °C before the experiment. Sequences and abbreviations are listed in Table 1 and Table S2. Thioflavin T (ThT) was ordered from Solarbio Life Science. Recombinant protein NONO (53-312) was synthesized by MerryBio Co.,Ltd. Recombinant protein DHX36 was purchased from OriGene Technologies Inc. RHAU53 peptide was synthesized by SynPeptide Co., Ltd.

**Table 1.**
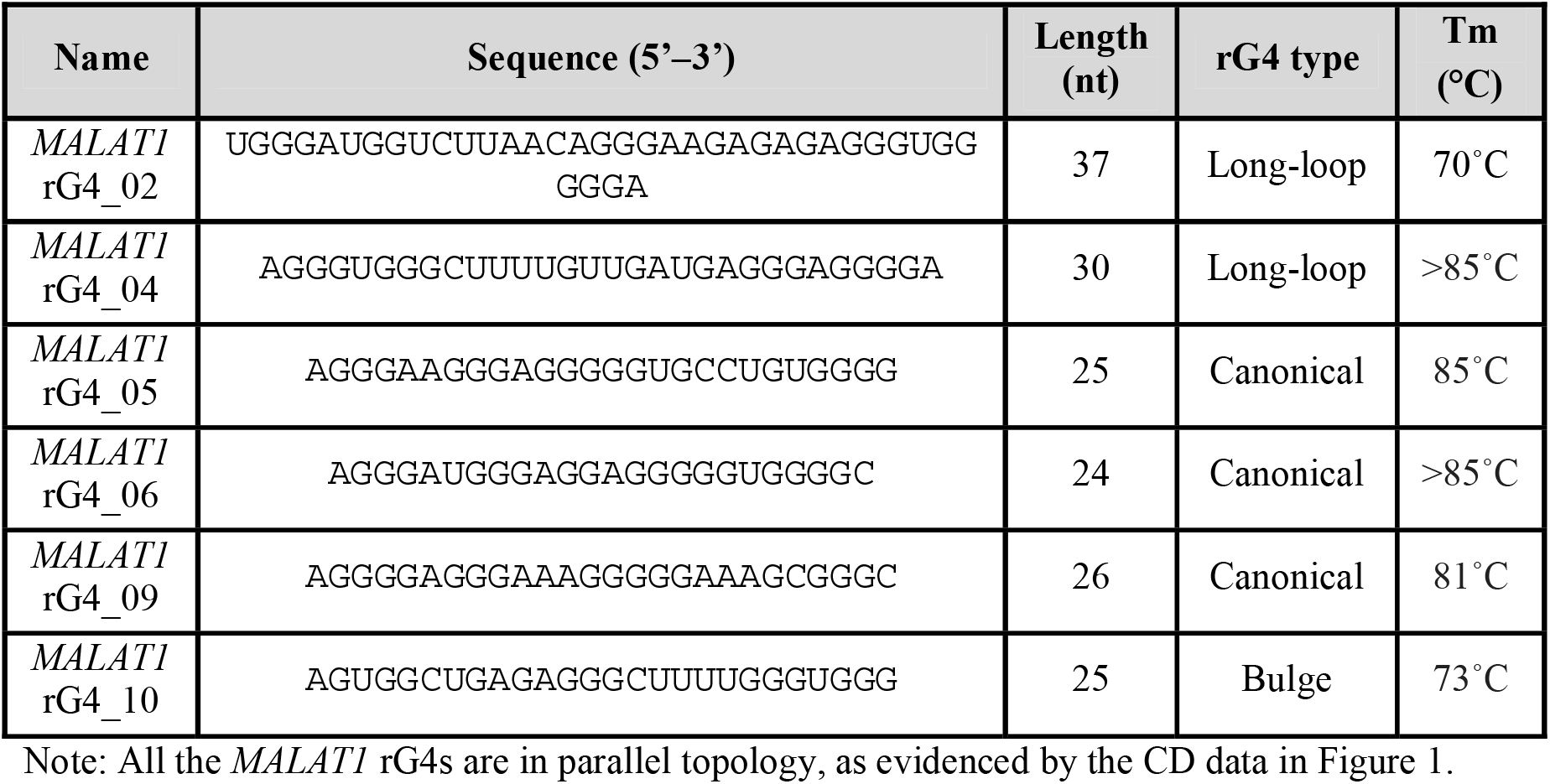
Oligonucleotide sequences of *MALAT1* rG4s used in this study.

### Circular Dichroism (CD) spectroscopy

CD assay was performed with a Jasco CD J-150 spectrometer and a 1 cm path length quartz cuvette. The reaction samples containing 5 μM oligos were prepared in 10 mM LiCac buffer (pH 7.0) and 150 mM KCl/LiCl to form total volume of 2 mL. The mixtures were then vortexed and heated at 95°C for 5 minutes and allowed to cool for 15 minutes at room temperature for renaturation. The samples were scanned at a 2 nm interval from 220 to 310 nm, and the data were blanked and normalised to mean residue ellipticity before smoothed over 10 nm. All the data were analyzed with Spectra Manager™ Suite and Microsoft Excel.

### UV melting assay

UV melting assay was performed with a Cary 100 UV-vis spectrophotometer and a 1 cm path length quartz cuvette. The reaction samples containing 5 μM oligos were prepared in 10 mM buffer (pH 7.0) and 150 mM KCl to form a total volume of 2 mL. The mixtures were vortexed and heated at 95°C for 5 minutes and allowed to cool for 15 minutes at room temperature for renaturation. In cuvettes sealed with Teflon tape, the samples were monitored at 295 nm from 5°C to 95°C with data collected over 0.5°C. The data were blanked and smoothed over 5 nm. All the data were analyzed with Microsoft Excel.

### Fluorescence spectroscopy

Fluorescence spectra was performed with a HORIBA FluoroMax-4 fluorescence spectrophotometer (Japan) and a 1 cm path length quartz cuvette. The reaction samples containing 1 μM oligos were prepared in prepared in 10 mM LiCac buffer (pH 7.0) and 150 mM KCl/LiCl to form a total volume of 100 μL. The mixtures were vortexed and heated at 95°C for 5 minutes and allowed to cool for 15 minutes at room temperature for renaturation. 1 μM of ThT ligand was added to the mixture. The emission spectra were collected from 440 to 700 nm with an excitation wavelength of 425 nm. The entrance and exit slits were 5 nm and 2 nm respectively, with the data collected every 2 nm. All the data were analyzed with Microsoft Excel.

### Comparative sequence analysis and gene conservation alignment

*MALAT1* gene sequences from different organisms were obtained from Ensembl Genome Browser (http://www.ensembl.org) using human *MALAT1* genes as reference. Multiple sequence alignments of the *MALAT1* genes were carried out with Jalview (52).

### G4 prediction analysis

G4 prediction analysis including cGcC (Consecutive G over consecutive C ratio) (53), G4H (G4Hunter) (54) and G4NN (G4 Neural Network) (55) was carried out by G4screener v.0.2 (http://scottgroup.med.usherbrooke.ca/G4RNA_screener/) with default setting. The G4 threshold is defined as cGcC > 4.5, G4H > 0.9, G4NN > 0.5.

### Electrophoretic Mobility Shift Assay (EMSA)

10 µl reaction mixtures containing 5 nM 5’ FAM-labeled RNA, 20 mM HEPES pH 7.5, 100 mM KCl, 0.12 mM EDTA, 25 mM Tris-HCl (pH 7.5), 10% glycerol and increasing concentrations of NONO(53-312) were prepared and incubated at 4°C for 1 hour. For inhibition assay, NONO(53-312) was increased to and fixed at 200 nM, and increasing concentrations of RHAU53 mixed with 5 nM 5’ FAM-labeled RNA and fixed amount of NONO(53-312). The RNA was heated at 75 □C for 5 minutes for denaturation before adding. The RNA-protein mixture were resolved by 4%, 37.5:1 (acrylamide: bis-acrylamide) non-denaturing polyacrylamide gel in 0.5 X Tris/Borate/EDTA (TBE) at 4°C, 80 V for 30 minutes (56). The gel was scanned by FujiFilm FLA-9000 Gel Imager at 650 V and quantified by ImageJ. The curve fitting and Kd/IC50 determination were carried out by Graphpad Prism using the one site-specific binding model.

### Filter-binding assay

50 µL reaction mixtures containing 2 nM 5’ Biotin-labelled RNA, 20 mM HEPES pH 7.5, 100 mM KCl, 0.12 mM EDTA, 25 mM Tris-HCl (pH 7.5) and increasing concentrations of NONO(53-312) were prepared and incubated at 4°C for 1 hour before loaded onto a Dot Blot apparatus (Bio-rad) containing a nitrocellulose membrane (top) and nylon membrane (bottom). For inhibition assay, Increasing concentrations of PDS/L-Apt.4-1c was mixed with 2 nM 5’ biotin-labeled RNA and fixed amount of NONO(53-312). The nylon membrane was rinsed with 0.5X TBE and nitrocellulose membrane was treated with 0.5 M KOH for 15 minutes at 4°C and rinse with 0.5X TBE before use. The membranes were wash 3 time with binding buffer before and after the mixture applied to the apparatus. The cross-linking step was performed under UV irradiation at 254 nm, 120,000 microjoules/cm^2^ for 5 minutes. The results were detected using a chemiluminescent nucleic acid detection module kit (Thermo Scientific) and scanned by ChemiDoc™ Touch Imaging System and quantified by ImageJ. The curve fitting and Kd determination were carried out by Graphpad Prism using the one site-specific binding model.

### Microscale thermophoresis (MST) binding assay

10 µL reaction mixtures containing 30 nM 5’ FAM-labeled RNA, 150 mM KCl, 1 mM MgCl_2_, 25 mM Tris-HCl (pH 7.5) and increasing concentrations of protein, ligand or aptamer were prepared and incubated at 37°C for 1 hour. For inhibition assay, Increasing concentrations of PDS/RHAU53/L-Apt.4-1c was mixed with 50 nM 5’ FAM-labeled RNA and fixed amount of NONO(53-312). The RNA and aptamer were heated at 75 □C for 5 minutes for denaturation before adding. The reaction mixtures were then loaded to MST capillary tubes (Nano-Temper Monolith NT.115), the measurement were carried out at 25°C using blue light mode and the binding affinity mode. The data was analysed by MST nano temper analysis (nta) analysis software. The curve fitting and Kd/IC50 determination were carried out by Graphpad Prism using the one site-specific binding model.

### RG4 pull-down assay

HEK293T (cell line authenticated and tested without mycoplasma contamination) nuclear lysate is extracted using NE-PER™ Nuclear extraction kit from Thermo Scientific™. Streptavidin C1 magnetic beads were blocked by 4 ng/ml yeast tRNA and 50 μg/ml BSA buffer for 30 minutes in room temperature and nuclear lysate were pre-cleared by incubating with 20 μL prewashed streptavidin C1 magnetic beads (Thermo Scientific™) for 1 hour in 4 °C. 300 pmol oligo and 60 μL prewashed streptavidin C1 magnetic beads were added for each reaction and the 5’ Biotin-labelled rG4 wildtype or mutant oligos were heated at 75 °C for 5 minutes for denaturation before adding to the mixture containing buffer A (10 mM Tris-HCl pH 7.5, 100 mM KCl, 0.1 mM EDTA, with/without 20 μM PDS). The beads immobilization step is processed in room temperature for 30 minutes in buffer A. The beads were washed for 3 times with buffer A. And 500 μg nuclear lysate was added in each reaction to the incubation mixture containing buffer B (20 mM Tris-HCl pH 7.5, 50 mM KCl, 0.5 mM EDTA, 10% glycerol). The incubation step is processed in 4°C for 2 hours and followed by 5 times stringent washes using buffer B. The enriched protein were eluted by heating at 95°C for 30 minutes in 1X Laemmli sample buffer (BioRad) and processed to western blot. The data was quantified by ImageJ and analyzed with Microsoft Excel.

### ChIRP-WB

300 million HEK293T cells were used per ChIRP-WB experiment. The cells were treated with 10 μM PDS (or DMSO as control) for 24 hours before collecting. The cells were crosslinked for 30 minutes in 3% formaldehyde, followed by 0.125 M glycine quenching for 5 minutes. The cells were wash by cold PBS and collected by centrifuge at 2000 rcf for 5 minutes. 100 mg cell pellets were dissolved in 1 mL cell lysis buffer (50 mM Tris-HCl pH 6.8, 1% SDS, 10 mM EDTA, 1 mM PMSF, protease inhibitor cocktail) and sonicated in 4°C using Bioruptor (Diagenode) until lysate is clear, centrifuged the cells at 18000 rcf at 4°C for 10 minutes and collected the supernatant. 10 μL of the control sample were saved as “input”. Each lysate sample was precleared with 30 μL streptavidin C1 magnetic beads (Thermo Scientific™). 2 mL hybridization buffer and 100 pmol probes per 1 mL of lysate were added to the mixture with 5’ Biotin-labelled *MALAT1* probes (Table S3) and rotated in 37 °C overnight. The hybridization buffer contains 750 mM NaCl, 50 mM Tris-HCl pH 6.8, 0.5% SDS, 1 mM EDTA, 15% formamide, 1 mM PMSF, protease inhibitor cocktail, RNase inhibitor. After the hybridization, 100 μL streptavidin C1 magnetic beads per 100 pmol probes were added and continue to mix samples at 37 °C for 30 minutes. The enriched RNA and RNA binding protein are isolated and processed to western blot and qPCR separately following the protocol (57). The data was analyzed with Microsoft Excel.

### Quantitative PCR (qPCR)

The RNA samples were resuspend in 100 µL proteinase K (pK) buffer containing 100 mM NaCl, 10 mM Tris–HCl pH 7.0, 1 mM EDTA, 0.5% SDS and 5% v/v pK and incubated in 50°C for 45 minutes and heated at 95 °C for 10 minutes (57). The RNA was isolated using Trizol:chloroform and ethanol and purified using RNeasy plus mini Kit (Qiagen). Reverse transcription was performed using PrimeScript™ RT Master Mix (TAKARA) and qPCR was carried out using SYBR Green Supermix (BioRad) and the SYBR program in CFX Connect Real-Time PCR Detection System. The primers used are listed in Table S2.

### Western blot

Samples for western blotting were resolved on 4-10% SDS-PAGE gels (BioRad) and transferred to immobilon-P PVDF membrane (Millipore). After blocking with 5% skim milk for 1 h at room temperature, the membrane was immunoblotted with primary antibodies diluted 1:1000 in blocking buffer for 1 hour at room temperature and washed 3 times with TBS-T for 15 minutes. Secondary antibodies were diluted 1:1000 in blocking buffer and applied for 1 hour at room temperature. The results were visualized using ChemiDoc™ Touch Imaging System and quantified by ImageJ. Primary antibodies used are as follows: Anti-nmt55 / p54nrb antibody (Abcam ab70335), Anti-GAPDH (Bio-station limited sc-32233).

## RESULTS

### Identification and characterization of G-quadruplex structures in *MALAT1* lncRNA

To explore rG4 in ncRNAs, we first examined our recently published *in vitro* rG4-seq dataset (5). While the rG4-seq dataset was collected using purified polyadenylated (polyA) RNA isolated from human HeLa cells, we have managed to identify more than 350 rG4s in a total of 203 ncRNA on the list (Table S1), such as lncRNAs *NEAT1* and *MALAT1*. We also employed the G4 prediction tools such as cGcC (53), G4hunter (54), and G4NN (55) on this ncRNA rG4 list, and found that 328 rG4 sequences showed at least with 1 score higher than the threshold set for those programs to score for G4, and 226 rG4 sequences showed all 3 scores above the threshold set (Table S1), suggesting the G4 prediction and rG4-seq data result are largely consistent. Using rG4-seq data and rG4 motif analysis, we have identified 15 and 11 rG4 forming sequences in lncRNA *NEAT1* and *MALAT1*, respectively (Table S1). Noteworthy, an independent study has recently predicted *NEAT1* to contain rG4 motifs and verified their formation using biophysical and biochemical assays (33), which further support our rG4-seq data. In this work, we focus on studying the formation of G-quadruplex structure in another important lncRNA *MALAT1*, and we selected 6 representative *MALAT1* rG4s according to the rG4 length and rG4 structural subtype. Overall, we selected 3 canonical rG4s and 3 non-canonical rG4s, including 2 long-loop rG4s and 1 bulge rG4, and the length are from 24 nt to 37 nt (Table 1).

To verify rG4 formation in *MALAT1* lncRNA, we have altogether carried out three spectroscopy assays. First, we conducted circular dichroism (CD) assay to determine the secondary structures and topologies of the 6 selected RNA oligonucleotides from *MALAT1* (Figure 1A-1F). The results showed that all 6 RNAs displayed a negative peak at 240 nm and a positive peak at 260 nm under physiologically-relevant K^+^ concentration (150 mM) (Figure 1A-1F), which suggest the formation of rG4 with parallel topology (58). Importantly, the CD signal is K^+^-dependent, and the CD signature of rG4 disappeared when the reaction was performed under the 150 mM Li^+^ conditions (Figure 1A-1F), supporting the distinctive CD profiles are caused by the formation of rG4 structure. Next, we performed UV melting assay under 150 mM K^+^ to verify rG4 formation to examine the thermostability of these 6 rG4s. Our data displayed a hypochromic shift at 295 nm, which is a hallmark of rG4 formation (Figure 1G-1L)(59). In addition, the UV melting results revealed that the melting temperature ™ of all 6 rG4 oligonucleotides are larger than 70 °C (Figure 1G-1L), which indicated that they are thermostable and folded into rG4 conformation under physiological temperature condition. Last, we carried out fluorescent turn-on spectroscopy assay using G-quadruplex-specific ligand thioflavin T (ThT)(60), and strong fluorescence enhancement was observed in 150 mM K^+^ when compared to 150 mM Li^+^ conditions for all 6 rG4 oligonucleotides (Figure 1M-1R), which further substantiate the formation of rG4 structures. Collectively, these spectroscopic assays verified that *MALAT1* lncRNA harbours several thermostable rG4 structures with parallel topology under physiological relevant potassium ion and temperature conditions.

**Figure 1.**
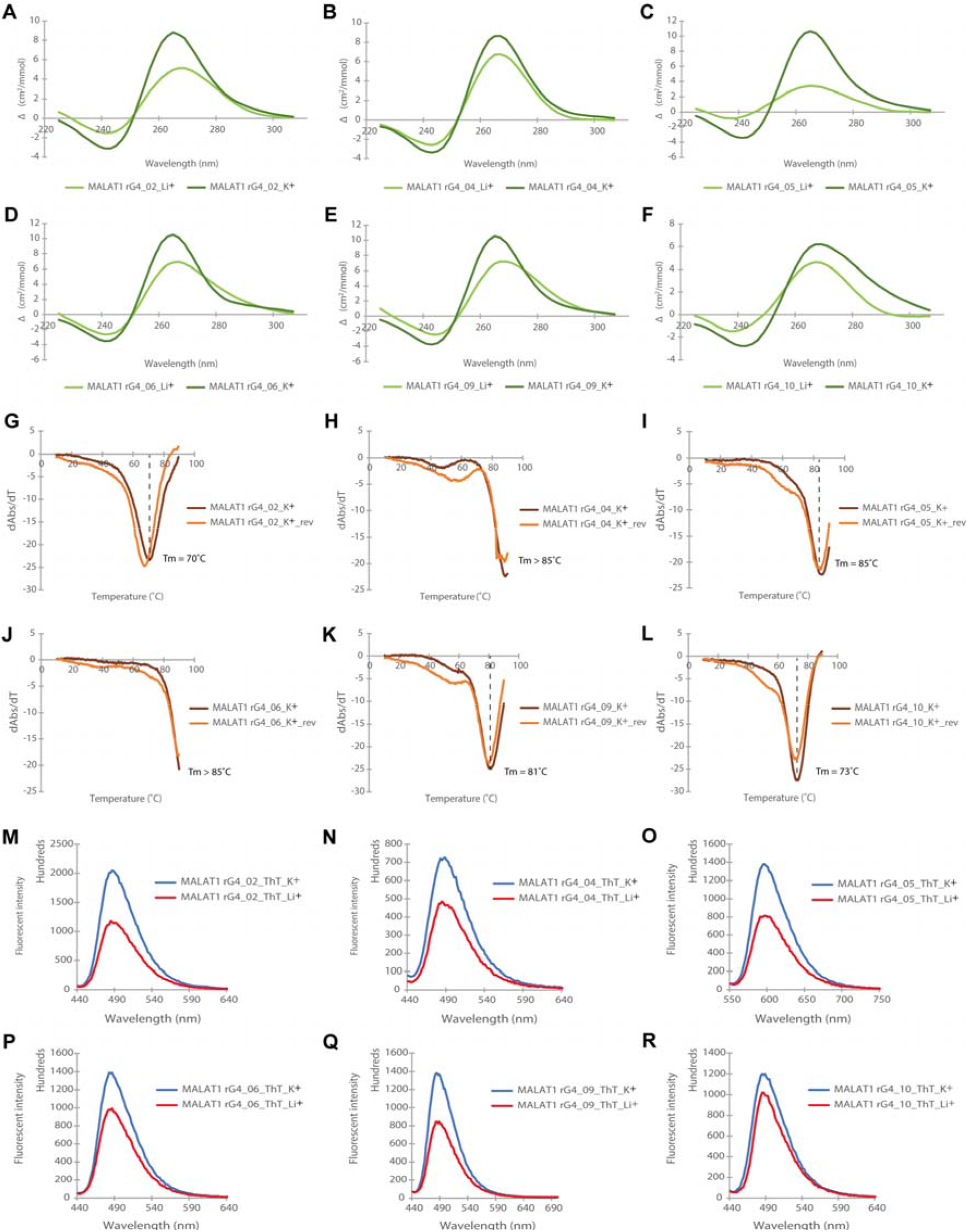
Biophysical characterization of *MALAT1* rG4 structures. (A)-(F) CD spectrum under K^+^ and Li^+^ conditions. The CD pattern becomes stronger in the presence of K^+^ rather than Li^+^, suggesting rG4 formation. The negative peak at 240 nm and the positive peak at 262 nm under K^+^ condition suggest a parallel topology of rG4. (G)-(L) UV melting. Hypochromic shift is observed at 295 nm, a strong sign of rG4 formation. The melting temperatures (Tms) for the *MALAT1* rG4s are indicated. (M)-(R) ThT enhanced fluorescence spectroscopy under K^+^ and Li^+^ conditions with an emission maximum wavelength of 490 nm. The fluorescence intensity of ThT is higher in rG4 under K^+^ compared to Li^+^, verifying the formation of rG4.

### *MALAT1* rG4s are conserved and interact with NONO protein *in vitro* and in nuclear lysate

Using comparative sequence analysis, we found that some of these *MALAT1* rG4s were highly conserved in 8 other species, including primates and mammals (Figure 2 and Figure S1). We also performed the G4 predictions on these rG4s and found they have largely similar scores as the human ones and passed the threshold set by the G4 prediction programs (Figure 2), suggesting these conserved *MALAT1* rG4s are likely to be formed and functional. Next, we attempted to identify proteins that can interact with the *MALAT1* rG4s. Studies have shown that RNA helicase DEAH box polypeptide 36 (DHX36) can bind rG4 with high affinity and selectivity (51,61), and it is found to localize both in the cytoplasm and nucleus (62). Therefore, we first performed binding assay with *MALAT1* rG4_04 and DHX36, and the result indicated that the *MALAT1* rG4_04 binds to DHX36 strongly with a dissociation constant (Kd) value of 130 ± 16 nM (Figure S2). Besides DHX36 protein, we hope to discover a new *MALAT1* rG4 binding protein in this study. *MALAT1* and *NEAT1* are both well conserved and nuclear-enriched lncRNAs. *MALAT1* lncRNA is localized in nuclear speckles, which are nuclear domains enriched in pre-mRNA splicing factors while *NEAT1* lncRNA is part of the paraspeckles, which are ribonucleoprotein bodies found in the interchromatin space of mammalian cell nuclei (63). Although *MALAT1* and *NEAT1* locate in different nuclear domains, they were found to localize adjacent to each other in the nucleus periodically and co-enrich many trans genomic binding sites and protein factors, such as NONO protein, indicating potential cooperation or complementary function between these two lncRNAs in regulating the formation of nuclear bodies (64). While NONO is one of the canonical paraspeckle proteins, it was also found in nuclear speckles, which suggest a diverse role of NONO (65,66). Recently, *NEAT1* was found to harbour several conserved rG4s which mediate the NONO-*NEAT1* association (33). To study the association of NONO protein with *MALAT1* rG4s, we first performed the electrophoretic mobility shift assay (EMSA) with purified recombinant protein NONO (53-312). NONO (53-312) contains two putative RNA-binding domain RRMs, the NONA/paraspeckle (NOPS) domain and half of the coiled-coil domain (33). The *NEAT1*_22619 works as a positive control to confirm that NONO (53-312) functions properly (Figure S3 and Table S2). We next tested the binding of *MALAT1* rG4s with NONO (53-312), and the EMSA results showed that NONO (53-312) exhibited strong binding to all 6 *MALAT1* rG4s, especially for *MALAT1* rG4_04 and *MALAT1* rG4_09 (Figure 3A-B and Figure S4) The Kd values were determined to be 46.9 ± 12.1 nM and 80.9 ± 18.1 nM respectively. To verify the binding is rG4-specific, we also designed two negative controls, a rG4 mutant, *MALAT1* rG4_04 MUT, and a non-G-quadruplex-forming scramble G-rich sequence (Table S2), and EMSA results showed both exhibited very weak binding to NONO (53-312) (Figure 3C-D). Besides, we also performed filter-binding assay with biotin-labelled *MALAT1* rG4_04 and *MALAT1* rG4_09, both wildtypes and mutant (Figure S5), and the results are consistent with EMSA (Figure 3E), illustrating *MALAT1* rG4 wildtypes, but not mutants, interact with NONO protein. Interestingly, we found that NONO (53-312) can compete with DHX36 and disrupt the binding between *MALAT1* rG4_04 and DHX36 (Figure S6).

**Figure 2.**
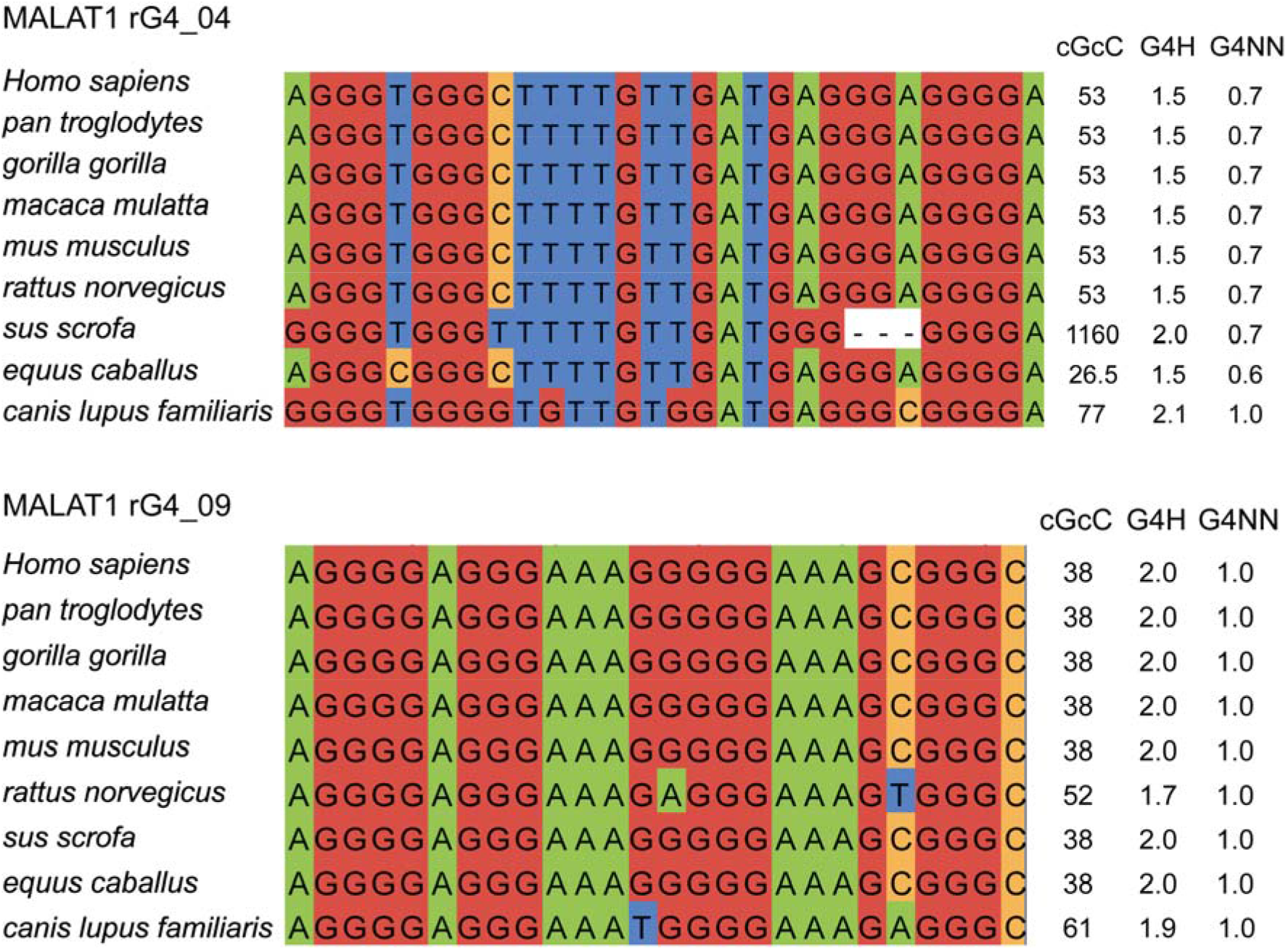
rG4 conservation analysis and predicted G4 scores for *MALAT1* rG4s. Comparative sequence analysis of *MALAT1* rG4_04 and *MALAT1* rG4_09 in different species and their predicted G4 scores. All G4 scores are higher than the threshold set in corresponding G4 prediction programs, i.e. cGcC > 4.5, G4H > 0.9, G4NN > 0.5, suggested the high likelihood of G4 formation. The Us in the rG4 sequences are replaced by Ts in the comparative sequence analysis and G4 prediction.

**Figure 3.**
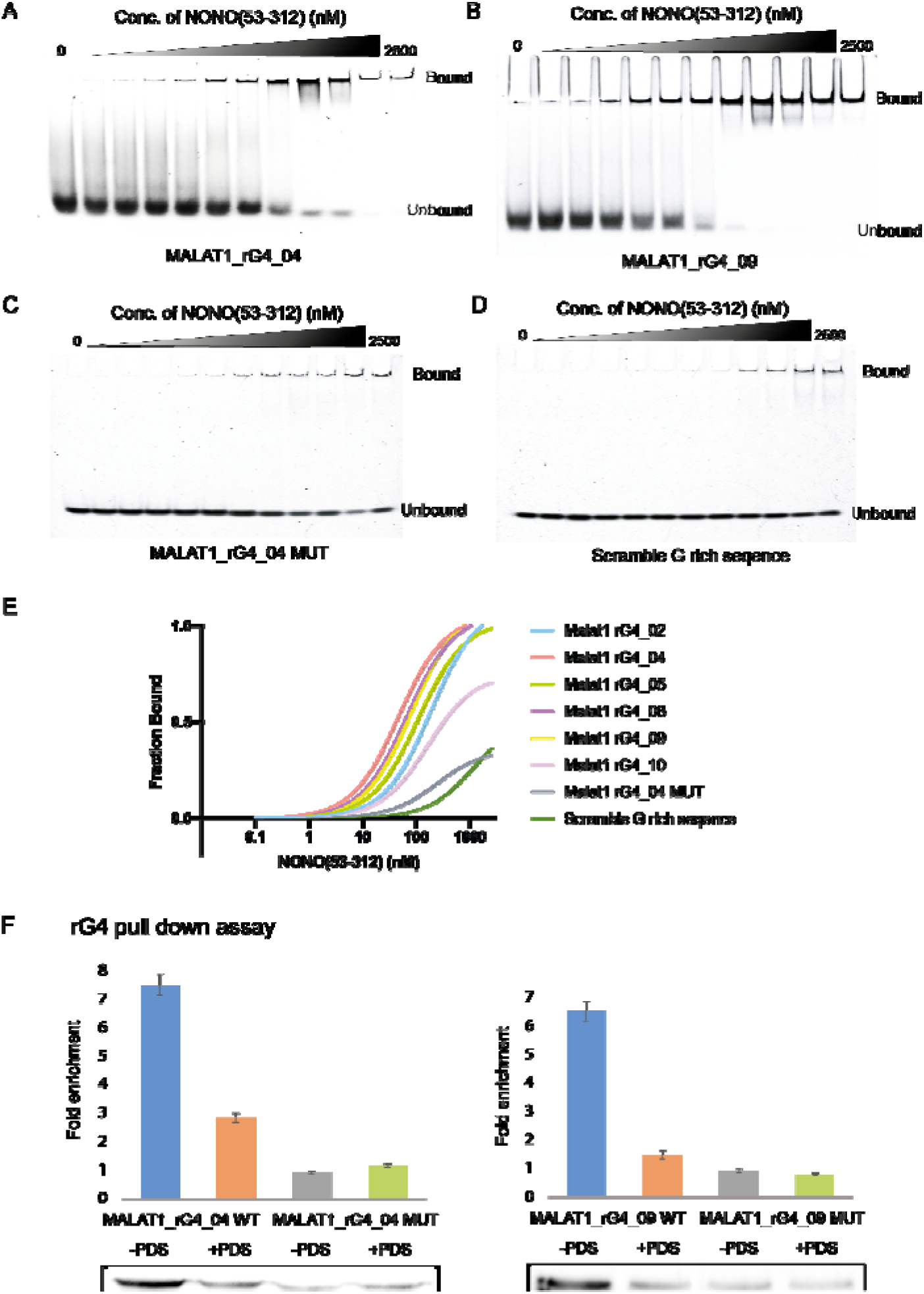
*MALAT1* rG4-NONO protein interactions *in vitro* and in nuclear lysate. (A) EMSA with recombinant NONO (53-312) and *MALAT1* rG4_04. (B) EMSA with recombinant NONO (53-312) and *MALAT1* rG4_09. (C) EMSA with recombinant NONO (53-312) and *MALAT1* rG4_04 MUT. (D) EMSA with recombinant NONO (53-312) and scramble G-rich sequence. (E) Binding curves of recombinant NONO (53-312) and *MALAT1* rG4_02, *MALAT1* rG4_04, *MALAT1* rG4_05, *MALAT1* rG4_06, *MALAT1* rG4_09, *MALAT1* rG4_10, *MALAT1* rG4_04 MUT and scramble G-rich sequence. The bindings of NONO to rG4s are stronger than rG4 mutant and scramble G-rich sequence, suggesting the binding is rG4-specific. (F) rG4 pull-down assay with HEK293T nuclear lysate. Western blot result of rG4 pulldown shows that NONO is enriched by *MALAT1*_rG4_04 and *MALAT1*_rG4_09 wildtype rG4s but not rG4 mutants, and the interaction can be disrupted by adding 20 μM PDS. In each blot, the rG4 mutant with no PDS treatment is normalized to one. Three independent experiments are performed. Error bars represent the S.E.M. The representative blot is shown here.

To strengthen the result and verify the interaction occur in more physiological-relevant settings, we next conducted pull-down experiments using biotin-labelled *MALAT1* rG4 oligos and attempted to pull down endogenous NONO protein from cell lysate obtained from HEK293T cells. Given that NONO protein and *MALAT1* lncRNA are both localized in nucleus, we carefully extracted the nuclear lysate using NE-PER™ Nuclear extraction kit. We used two pairs of RNA oligonucleotides, *MALAT1* rG4_04 WT and *MALAT1*_rG4_04 MUT, as well as *MALAT1*_rG4_09_WT and *MALAT1*_rG4_09_MUT, and showed that under normal conditions, the wildtype versions (both *MALAT1* rG4_04 WT and *MALAT1*_rG4_09_WT) were able to pull down endogenous NONO protein more than the mutant version (*MALAT1* rG4_04 MUT and *MALAT1*_rG4_09_MUT), with a ratio of 9:1 and 7:1, respectively (Figure 3F). G4 ligands, such as pyridostatin (PDS), have been previously reported to inhibit and display native G4-protein interaction (67,68). To support the binding relies on *MALAT1* rG4 structure, PDS was added to the reaction to a final concentration of 20 μM, and a drop in the ratio was observed, suggesting PDS binds to *MALAT1* rG4 and displace the rG4-protein interaction. Altogether, the above results provided clear evidence that *MALAT1* rG4s can interact with NONO protein *in vitro* and in nuclear cell lysate.

### *MALAT1* interacts with NONO via rG4 structure in cells

Having found that NONO recognizes *MALAT1* rG4s both *in vitro* and in cell lysate, we aspired to ascertain whether the preference for *MALAT1* rG4s detected is reflected in living cells as well. To examine the binding between *MALAT1* and NONO in cells, we first designed and applied a total of 40 biotinylated *MALAT1* probes to carry out an RNA binding protein detection method referred to as ChIRP-western blot (ChIRP-WB) (69) (Table S3). Our rationale is that if NONO protein interacts with *MALAT1* lncRNA in cells, then after crosslinking, sonication, biotinylated *MALAT1* lncRNA probe capture and enrichment, the NONO protein should be enriched in the western blot assay (Figure 4A). Besides, in order to determine whether the rG4 structures in *MALAT1* participate in the *MALAT1*-NONO interaction in cells, we also treated another set of HEK293T cells with 10 μM PDS (or DMSO as control) for 24 hours before crosslinking (Figure 4A). We also conducted negative control with RNase A treatment. Compared with the negative control with RNase treatment, NONO was significantly enriched in both groups (with/without PDS), which demonstrated that *MALAT1* interacts with NONO in cells. Importantly, we found that the *MALAT1*-NONO interaction was weakened by adding G4-specific ligand PDS, indicating NONO protein recognizes *MALAT1* lncRNA through rG4 structures. This is also in agreement with our observation in the pull-down assay, where the addition of PDS dissociated the *MALAT1* rG4-NONO protein binding in nuclear lysate (Figure 3). GAPDH protein, which is not known to bind to *MALAT1*, was used as a negative control for ChIRP-WB experiments, and no or very weak band can be detected under all 3 conditions tested (Figure 4B). Furthermore, to validate the specificity and the efficiency of biotinylated *MALAT1* probes, we carried out qPCR using 2 sets of *MALAT1* primers and 1 set of *Xist* primers (Table S2). The results showed that *MALAT1* was greatly enriched while *Xist* was not, highlighting that the biotinylated *MALAT1* probes can capture and enrich *MALAT1* lncRNA successfully and efficiently (Figure 4C). Combining both *in vitro*, in nuclear lysate (Figure 3), and in cell data here (Figure 4), we have provided substantial evidence that *MALAT1* rG4s can strongly interact with NONO both *in vitro*, in cell lysate, and in cells.

**Figure 4.**
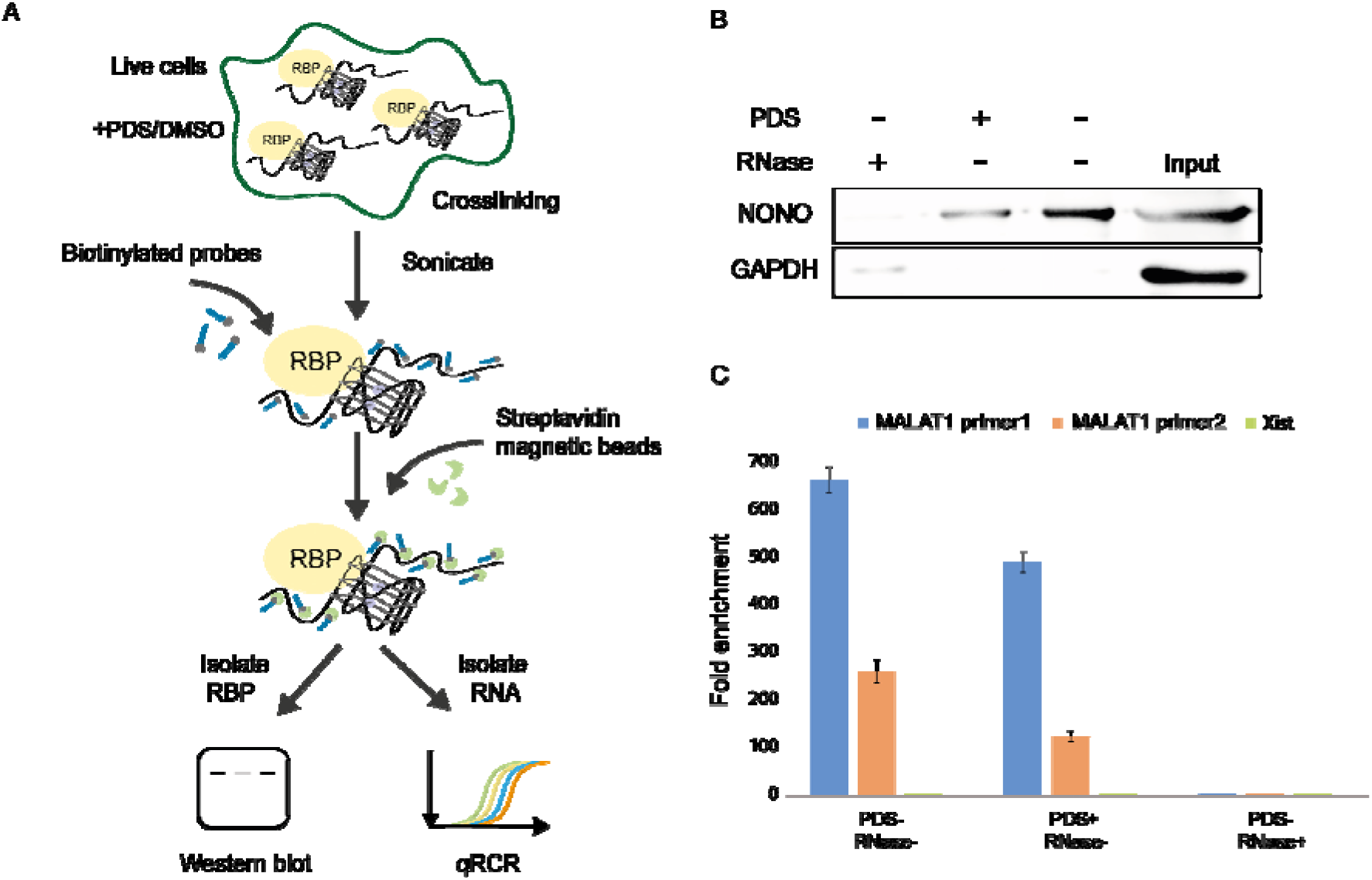
*MALAT1* rG4-NONO protein interaction in cells. (A) Scheme for the ChIRP-WB and qPCR. RNA binding protein (RBP) is crosslinked to lncRNA of interest *in vivo*, followed by sonication. Biotinylated tiling probes are then hybridized to target lncRNA, and the complexes are purified using magnetic streptavidin beads, followed by stringent washes. The enriched lncRNA and RBP are isolated and processed by western blot and qPCR separately. (B) NONO protein is enriched by *MALAT1* ChIRP, and their interaction can be disrupted by 10 μM G4 ligand PDS, suggesting the binding is mediated by rG4. GAPDH is used as negative control in this experiment. Three independent experiments are performed. (C) *MALAT1* RNA is pulled down by biotinylated probes, and PDS treatment do not have much effect on the pull-down efficiency. Both sets of *MALAT1* qPCR primers shows great *MALAT1* lncRNA enrichment. *Xist* lncRNA is used as a negative control in this experiment and no enrichment is observed. Three independent experiments are performed. Error bars represent the S.E.M.

### *MALAT1* rG4-NONO interaction can be suppressed by multiple G4 targeting tools

Motivated by the finding that *MALAT1* lncRNA interacts with NONO via rG4 structure, we wonder if the interaction between *MALAT1* rG4 and NONO can be disrupted by G4 targeting tools such as G4-specific small molecule, peptide, and aptamer (Figure 5A). Currently, G4 targeting with small molecule is the most commonly employed strategy to recognize G4 motifs specifically and interfere with G4–protein interactions (70). We first tested whether *MALAT1* rG4_04 can interact with G4-specific ligand PDS using MST and found the G4-PDS complex to form at around 20 nM PDS, with a Kd value of 66 ± 8 nM (Figure S7). To determine the ability of PDS to inhibit *MALAT1* rG4–NONO interaction, we assembled the *MALAT1* rG4_04–NONO (53-312) complex and titrated it with an increasing concentration of PDS (0-1000 nM). The MST result showed that PDS could effectively suppress *MALAT1* rG4_04–NONO (53-312) interaction with a half-maximal inhibitory concentration (IC50) value of 498.2 ± 94.5 nM (Figure 5B), suggesting that PDS can successfully dissociate *MALAT1* rG4_04–NONO (53-312) interaction even when the concentration of NONO (53-312) is as high as 200 nM. Noteworthy, this result further supports our above-mentioned data that PDS can interfere with the interaction between *MALAT1* lncRNA and NONO in nuclear lysate and in cell (Figures 3 and 4). Next, we assessed the interference of *MALAT1* rG4_04–NONO (53-312) interaction using the G4-specific peptide approach. As mentioned above, DHX36, or RHAU, is a family member of the ATP-dependent RNA helicase that specifically binds to and resolves parallel-stranded G-quadruplexes (71). Recently, the key protein region involving in G4 interactions were determined to be near the N-terminus of RHAU protein, and one of the widely used and studied is RHAU53, which contains 53-amino acids long RHAU protein fragment (72). As such, we used RHAU53 peptide as a G4 targeting peptide in this inhibition assay (Table S2). Similar as PDS, we constructed the *MALAT1* rG4_04–NONO (53-312) complex followed by an escalating concentration of RHAU53. From the MST data, the IC50 was determined to be 621.1 ± 95.3 nM (Figure 5C), which is comparable with the PDS result. Moreover, we demonstrated the *MALAT1* rG4_04–NONO (53-312) interaction can be suppressed by rG4-targeting L-RNA aptamer, which is a new class of G4-targeting tool. L-Apt.4-1c is a novel L-RNA aptamer developed by our group recently (73). It was reported to generally bind to rG4s and interfere with rG4–protein interaction (73,74). In this study, we first tested the binding affinity between *MALAT1* rG4_04 and L-Apt.4-1c and found the Kd to be 165 ± 45 nM (Figure S8), indicating they strongly and directly interact with each other. To assess its capability of suppressing *MALAT1* rG4_04–NONO (53-312) interaction, we performed MST inhibition assay and monitored the IC50 to be 618.1 ± 110.7 nM (Figure 5D), suggesting the inhibition ability of L-Apt.4-1c is analogous with PDS and RHAU53.

**Figure 5.**
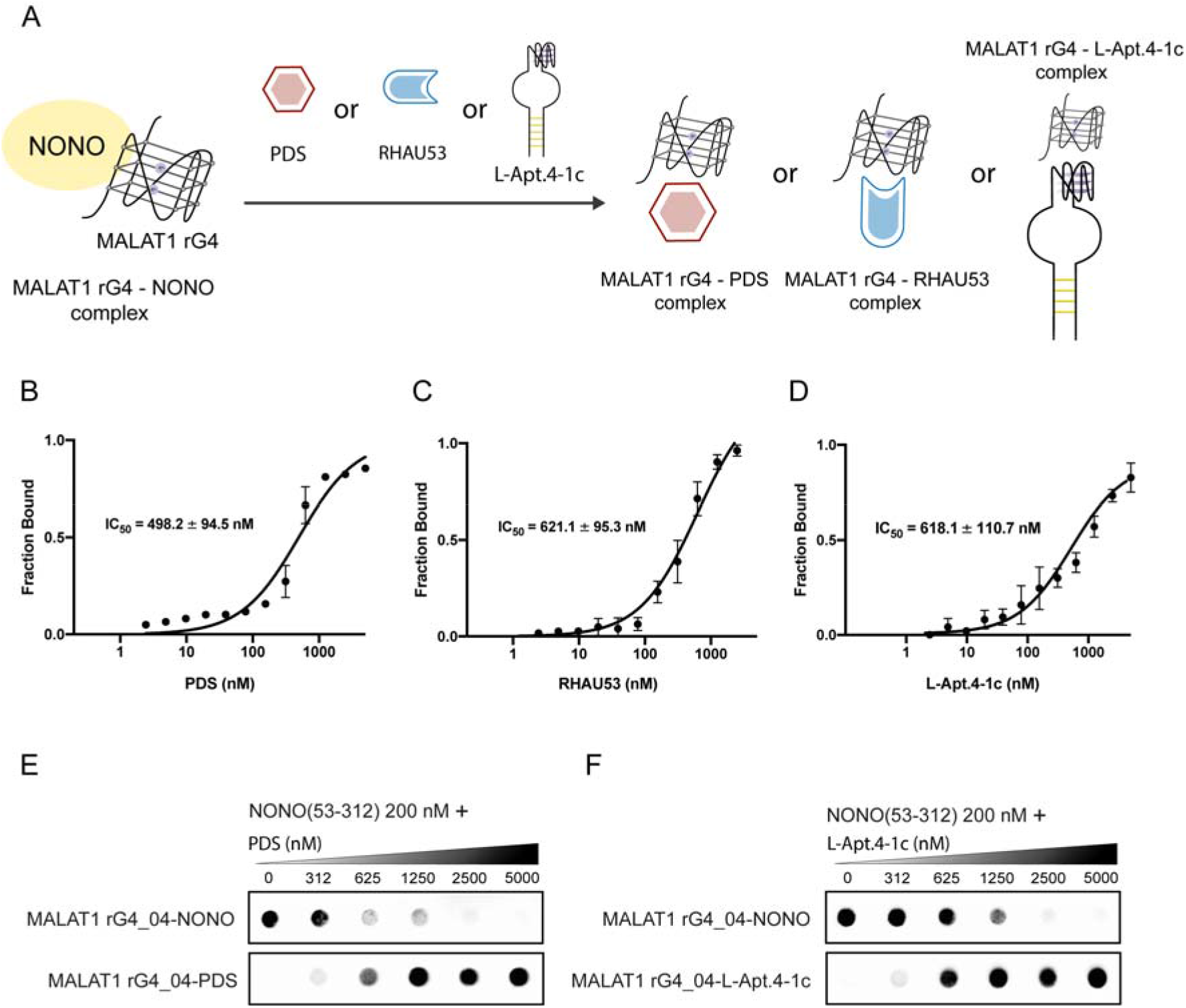
Suppression of *MALAT1* rG4-NONO interaction by multiple G4 targeting tools. (A) Schematic representation of *MALAT1* rG4-NONO interaction that is interfered by various G4 targeting tools including small molecule, peptide, and L-aptamer. (B) Saturation plot of PDS for its inhibition of *MALAT1* rG4-NONO interaction. Reaction mixture contains 50 nM FAM *MALAT1* rG4_04, 200 nM NONO (53-312) and increasing concentrations of PDS (0.15–5000 nM). The IC50 is found to 498.2 ± 94.5 nM. (C) Similar set up as (B) except RHAU53 is used. The IC50 value is found to be 621.1 ± 95.3 nM. (D) Similar set up as (B) except L-Apt.4-1c was used. The IC50 value is found to be 618.1 ± 110.7 nM. (E) Filter-binding result of PDS for its inhibition of *MALAT1* rG4-NONO interaction. Reaction mixture contains 2 nM Biotin *MALAT1* rG4_04, 150 nM NONO (53-312) and increasing concentrations of PDS (0.15–5000 nM). (F) Similar set up as (E) except L-Apt.4-1c is used.

To support the MST results above, we carried out two independent assays, filter-binding assay and EMSA, and the results were consistent with our MST data. In filter-binding assay, by employing the reaction mixture to a Dot Blot apparatus containing a nitrocellulose membrane (top) and nylon membrane (bottom), rG4-protein bound would retain in the top membrane while the bottom membrane detained the rest. From the data, the *MALAT1* rG4_04–NONO (53-312) complex was dissociated when increasing the concentration of PDS or L-Apt.4-1c (Figure 5E-F). Our EMSA result showed that as the concentration of RHAU53 escalates from 0 to 2500 nM, the *MALAT1* rG4_04–NONO (53-312) complex band declined while the *MALAT1* rG4_04–RHAU53 complex band increased (Figure S9), suggesting the interference of RHAU53 to *MALAT1* rG4_04–NONO (53-312) interaction. Taken the results together, we have demonstrated that the interaction between *MALAT1* rG4 and NONO can be suppressed by multiple G4 targeting tools including G4-specific small molecule, peptide, and L-RNA aptamer, which provide diverse approaches to targeting *MALAT1* and its RNA-protein complex via its rG4 motifs.

## DISCUSSION

Recent studies have shown that lncRNAs play vital roles in different biological processes (75-77). However, few works have elucidated RNA structural elements in lncRNAs (78-81). rG4s, being non-classical RNA structural motifs, have been studied more extensively in mRNAs, but less so in ncRNAs such as lncRNAs (13,14,82). Besides the well-characterized rG4s in human telomerase RNA (*hTR*) (30,51,83) and Telomeric repeat-containing RNA (*TERRA*) (31,32,84), there is so far only a handful of rG4 reported in lncRNAs, including *FLJ39051*(*GSEC)* (29), *LUCAT1* (49), and *NEAT1* (33). LncRNA *MALAT1* is a highly conserved cancer-associated lncRNA that interacts with RNA-binding proteins to participate in transcription and post-transcriptional regulation in cells (85,86). It promotes cell proliferation, metastasis, and invasion *in vitro* and *in vivo*, thus playing a critical role in regulating cancer progression and metastasis (85-87). In this work, we have provided the first evidence that lncRNA *MALAT1* contains multiple rG4 structures. By applying different biophysical and biochemical assays, we demonstrated that these rG4s are thermostable and formed in physiologically relevant conditions (Figure 1). Moreover, using comparative sequence analysis, we found *MALAT1* rG4s that we study in this work are conserved, and some of them are highly conserved across 8 other species (Figure 2 and Figure S1), including primates and mammals. Importantly, the G4 prediction scores were all above the G4 threshold (Figure 2), indicating these sequences are likely to form rG4 structure. Furthermore, the conservation of the rG4 structure hints at the potential significance of the G4 structural element in the function of *MALAT1*.

*MALAT1* and *NEAT1* are both abundant and nuclear-enriched lncRNAs that localize adjacent to each other in the nucleus periodically and co-enrich many trans-genomic binding sites and protein factors, including NONO protein (64-66). The affinity of NONO for rG4 structures has been observed very recently for *NEAT1* rG4s (33). In this study, we have found for the first time that NONO interacts with *MALAT1* via rG4 structures both *in vitro* and *in vivo* (Figure 3-4). We hypothesize that the rG4 structure motif may serve the role of potential cooperation or complementary function between these two lncRNAs in recruiting NONO and other nuclear proteins and regulating the formation of nuclear bodies. Our speculation is also supported by data that suggested NONO is also located in both paraspeckles and nuclear speckles (65,88). In addition, SFPQ (Splicing factor proline- and glutamine-rich), a binding partner of NONO, was also reported to interact with *NEAT1* rG4 (33) and *MALAT1* (85,86), which further supports our hypothesis. It is also possible that rG4 helicase such as DHX36 may participate in the regulation of rG4-NONO complex formation. Our data showed that it binds to *MALAT1* rG4 strongly, and DHX36 and NONO protein can compete for *MALAT1* rG4 (Figure S2 and S6). As such, the interaction between *MALAT1* rG4 and NONO, as well as DHX36, may be a promising target for *MALAT1/NEAT1-*related diseases and cancer therapy.

To explore the potential application, we have investigated the disruption ability of three classes of G4 targeting tools, including G4-specific small molecule, peptide and L-RNA aptamer. PDS is one of the popular G4 ligands used in recognizing G4 motifs and interfering with the G4–protein interactions (67,68). In this work, we have utilized PDS for both in vitro, in nuclear cell lysate, and in cell assays, underscoring its versatility and application in studying rG4 in lncRNA. RHAU53 peptide is an emerging G4 targeting tool that can specifically identify parallel G4, and a number of improved versions have been recently developed for specific G4 targeting (72,89-91). rG4-targeting L-RNA aptamer is a new class of G4-targeting tool introduced by us (56). L-Apt.4-1c is a novel L-RNA aptamer developed by our group recently (73). It was reported to generally bind to rG4s and interfere with the rG4–protein interaction *in vitro* and in cell lysate (73,74). Notably, L-Apt.4-1c is the shortest L-aptamer developed so far, with only 25 nt in length, making it easier to apply to *in vivo* experiments in the near future. Based on our data, L-Apt.4-1c exhibited comparable ability to disrupt *MALAT1* rG4-NONO interaction compared to PDS and RHAU53, highlighting the potential of L-aptamer in the G4 targeting field and the possibility of targeting lncRNA rG4-protein interactions. In addition, modification of the L-Apt.4-1c, such as cyclization (74), and optimization of the L-aptamer sequence should improve its performance further. It will be interesting and one of our future directions to apply these G4 targeting tools to explore the potential regulatory and mechanistic role of rG4s in *MALAT1* lncRNA-associated cancers.

As described early in the result section, we have re-analyzed and identified more than 350 rG4 in about 200 ncRNAs using our published rG4-seq dataset. In this list (Table S1), we also found rG4s in other biologically relevant lncRNAs such as *TUG1* (taurine upregulated gene 1), *HCG11* (HLA Complex Group 11), *DGCR5* (DiGeorge Syndrome Critical Region Gene 5), *HOTAIRM1* (HOXA Transcript Antisense RNA, Myeloid-Specific 1), and many more, which have been functionally characterized in other studies recently (92-99). Our view is that this list is likely an underestimation of the rG4s in ncRNAs as the rG4-seq was polyA-enriched (5), and many lncRNAs do not have polyA tail or polyA region, which will be omitted in the dataset. However, we have clearly illustrated in this study that the rG4 candidates obtained in this list are largely consistent with the G4 prediction tools results, suggested that a future computational approach to look for lncRNA rG4s or even all ncRNAs rG4 is likely possible and will yield valuable insights for experimentalists. Notably, based on our findings in this study on the rG4s in *MALAT1* lncRNA, it is promising to apply the strategies and methods reported here to investigate the rG4 structures and protein interactions in other important lncRNAs *in vitro* and *in vivo*.

## CONCLUSION

In sum, we have revealed novel rG4s in *MALAT1* lncRNAs for the first time, and showed that these rG4s are thermostable and conserved. We also uncovered these rG4s to have specific interactions with NONO protein both *in vitro*, in nuclear lysate and in cells. Importantly, we demonstrated that rG4 targeting tools including small molecule, peptide, and L-RNA aptamer can be used to suppress the *MALAT1* rG4-NONO protein interaction, providing diverse strategies to targeting *MALAT1* and its RNA-protein complex via its rG4 motifs. The findings reported in this study will motivate further exploration of the role of rG4s in *MALAT1* lncRNA biology, and the approaches presented here should be generally applicable to the study of rG4 in other functional lncRNAs.

## SUPPLEMENTARY DATA

Supplementary Data are available online.

## DATA AVAILABILITY

The data generated during all experiments is available from the author upon reasonable request.

## ACKNOWLEDGEMENT

We thank Dr. Omer Ziv, Eugene Yui-Ching Chow, Jia-Hao Yuan and Haizhou Zhao for the discussion and inputs in this manuscript.

## REFERENCES

1. Neidle, S. and Balasubramanian, S. (2006) Quadruplex Nucleic Acids, Vol. 7. Royal Society of Chemistry. Cambridge, UK.

2. Kwok, C.K. and Merrick, C.J. (2017) G-Quadruplexes: Prediction, Characterization, and Biological Application. Trends in Biotechnology, 35, 997–1013.

3. Huppert, J.L. and Balasubramanian, S. (2005) Prevalence of quadruplexes in the human genome. Nucleic Acids Res, 33, 2908–2916.

4. Chambers, V.S., Marsico, G., Boutell, J.M., Di Antonio, M., Smith, G.P. and Balasubramanian, S. (2015) High-throughput sequencing of DNA G-quadruplex structures in the human genome. Nat. Biotechnol., 33, 877–881.

5. Kwok, C.K., Marsico, G., Sahakyan, A.B., Chambers, V.S. and Balasubramanian, S. (2016) rG4-seq reveals widespread formation of G-quadruplex structures in the human transcriptome. Nat. Methods, 13, 841–844.

6. Lightfoot, H.L., Hagen, T., Tatum, N.J. and Hall, J. (2019) The diverse structural landscape of quadruplexes. FEBS Lett, 593, 2083–2102.

7. Banco, M. and Ferre-D’Amare, A. (2021) The emerging structural complexity of G-quadruplex RNAs. RNA.

8. Chan, C.Y., Umar, M.I. and Kwok, C.K. (2019) Spectroscopic analysis reveals the effect of a single nucleotide bulge on G-quadruplex structures. Chem Commun (Camb), 55, 2616–2619.

9. Kwok, C.K., Sherlock, M.E. and Bevilacqua, P.C. (2013) Effect of loop sequence and loop length on the intrinsic fluorescence of G-quadruplexes. Biochemistry, 52, 3019–3021.

10. Spiegel, J., Adhikari, S. and Balasubramanian, S. (2020) The Structure and Function of DNA G-Quadruplexes. Trends Chem, 2, 123–136.

11. Varshney, D., Spiegel, J., Zyner, K., Tannahill, D. and Balasubramanian, S. (2020) The regulation and functions of DNA and RNA G-quadruplexes. Nat Rev Mol Cell Biol.

12. Robinson, J., Raguseo, F., Nuccio, S.P., Liano, D. and Di Antonio, M. (2021) DNA G-quadruplex structures: more than simple roadblocks to transcription? Nucleic Acids Res.

13. Tassinari, M., Richter, S.N. and Gandellini, P. (2021) Biological relevance and therapeutic potential of G-quadruplex structures in the human noncoding transcriptome. Nucleic Acids Res, 49, 3617–3633.

14. Lyu, K., Chow, E.Y., Mou, X., Chan, T.F. and Kwok, C.K. (2021) RNA G-quadruplexes (rG4s): genomics and biological functions. Nucleic Acids Res, 49, 5426–5450.

15. Rouleau, S., Glouzon, J.S., Brumwell, A., Bisaillon, M. and Perreault, J.P. (2017) 3’ UTR G-quadruplexes regulate miRNA binding. RNA, 23, 1172–1179.

16. Stefanovic, S., Bassell, G.J. and Mihailescu, M.R. (2015) G quadruplex RNA structures in PSD-95 mRNA: potential regulators of miR-125a seed binding site accessibility. RNA, 21, 48–60.

17. Subramanian, M., Rage, F., Tabet, R., Flatter, E., Mandel, J.L. and Moine, H. (2011) G-quadruplex RNA structure as a signal for neurite mRNA targeting. Embo Reports, 12, 697–704.

18. Goering, R., Hudish, L.I., Guzman, B.B., Raj, N., Bassell, G.J., Russ, H.A., Dominguez, D. and Taliaferro, J.M. (2020) FMRP promotes RNA localization to neuronal projections through interactions between its RGG domain and G-quadruplex RNA sequences. Elife, 9.

19. Crenshaw, E., Leung, B.P., Kwok, C.K., Sharoni, M., Olson, K., Sebastian, N.P., Ansaloni, S., Schweitzer-Stenner, R., Akins, M.R., Bevilacqua, P.C. et al.. (2015) Amyloid Precursor Protein Translation Is Regulated by a 3’UTR Guanine Quadruplex. PLoS One, 10, e0143160.

20. Murat, P., Marsico, G., Herdy, B., Ghanbarian, A.T., Portella, G. and Balasubramanian, S. (2018) RNA G-quadruplexes at upstream open reading frames cause DHX36- and DHX9-dependent translation of human mRNAs. Genome Biol, 19, 229.

21. Beaudoin, J.D. and Perreault, J.P. (2013) Exploring mRNA 3’-UTR G-quadruplexes: evidence of roles in both alternative polyadenylation and mRNA shortening. Nucleic Acids Res, 41, 5898–5911.

22. Marcel, V., Tran, P.L., Sagne, C., Martel-Planche, G., Vaslin, L., Teulade-Fichou, M.P., Hall, J., Mergny, J.L., Hainaut, P. and Van Dyck, E. (2011) G-quadruplex structures in TP53 intron 3: role in alternative splicing and in production of p53 mRNA isoforms. Carcinogenesis, 32, 271–278.

23. Westmark, C.J. and Malter, J.S. (2007) FMRP mediates mGluR(5)-dependent translation of amyloid precursor protein. Plos Biology, 5, 629–639.

24. Thandapani, P., Song, J., Gandin, V., Cai, Y., Rouleau, S.G., Garant, J.M., Boisvert, F.M., Yu, Z., Perreault, J.P., Topisirovic, I. et al.. (2015) Aven recognition of RNA G-quadruplexes regulates translation of the mixed lineage leukemia protooncogenes. Elife, 4.

25. Huang, H., Zhang, J., Harvey, S.E., Hu, X. and Cheng, C. (2017) RNA G-quadruplex secondary structure promotes alternative splicing via the RNA-binding protein hnRNPF. Genes Dev, 31, 2296–2309.

26. Shahid, R., Bugaut, A. and Balasubramanian, S. (2010) The BCL-2 5’ untranslated region contains an RNA G-quadruplex-forming motif that modulates protein expression. Biochemistry, 49, 8300–8306.

27. Beaudoin, J.D. and Perreault, J.P. (2010) 5’-UTR G-quadruplex structures acting as translational repressors. Nucleic Acids Research, 38, 7022–7036.

28. Mattick, J.S. and Makunin, I.V. (2006) Non-coding RNA. Hum Mol Genet, 15 Spec No 1, R17–29.

29. Matsumura, K., Kawasaki, Y., Miyamoto, M., Kamoshida, Y., Nakamura, J., Negishi, L., Suda, S. and Akiyama, T. (2017) The novel G-quadruplex-containing long non-coding RNA GSEC antagonizes DHX36 and modulates colon cancer cell migration. Oncogene, 36, 1191–1199.

30. Lattmann, S., Stadler, M.B., Vaughn, J.P., Akman, S.A. and Nagamine, Y. (2011) The DEAH-box RNA helicase RHAU binds an intramolecular RNA G-quadruplex in TERC and associates with telomerase holoenzyme. Nucleic Acids Res, 39, 9390–9404.

31. Biffi, G., Tannahill, D. and Balasubramanian, S. (2012) An intramolecular G-quadruplex structure is required for binding of telomeric repeat-containing RNA to the telomeric protein TRF2. J. Am. Chem. Soc., 134, 11974–11976.

32. Deng, Z., Norseen, J., Wiedmer, A., Riethman, H. and Lieberman, P.M. (2009) TERRA RNA binding to TRF2 facilitates heterochromatin formation and ORC recruitment at telomeres. Mol Cell, 35, 403–413.

33. Simko, E.A.J., Liu, H., Zhang, T., Velasquez, A., Teli, S., Haeusler, A.R. and Wang, J. (2020) G-quadruplexes offer a conserved structural motif for NONO recruitment to NEAT1 architectural lncRNA. Nucleic Acids Res, 48, 7421–7438.

34. Chan, K.L., Peng, B., Umar, M.I., Chan, C.Y., Sahakyan, A.B., Le, M.T.N. and Kwok, C.K. (2018) Structural analysis reveals the formation and role of RNA G-quadruplex structures in human mature microRNAs. Chem. Commun. (Camb), 54, 10878–10881.

35. Kwok, C.K., Sahakyan, A.B. and Balasubramanian, S. (2016) Structural Analysis using SHALiPE to Reveal RNA G-Quadruplex Formation in Human Precursor MicroRNA. Angew Chem Int Ed Engl, 55, 8958–8961.

36. Pandey, S., Agarwala, P., Jayaraj, G.G., Gargallo, R. and Maiti, S. (2015) The RNA Stem-Loop to G-Quadruplex Equilibrium Controls Mature MicroRNA Production inside the Cell. Biochemistry, 54, 7067–7078.

37. Vourekas, A., Zheng, K., Fu, Q., Maragkakis, M., Alexiou, P., Ma, J., Pillai, R.S., Mourelatos, Z. and Wang, P.J. (2015) The RNA helicase MOV10L1 binds piRNA precursors to initiate piRNA processing. Genes Dev, 29, 617–629.

38. Balaratnam, S., Hettiarachchilage, M., West, N., Piontkivska, H. and Basu, S. (2019) A secondary structure within a human piRNA modulates its functionality. Biochimie, 157, 72–80.

39. Mestre-Fos, S., Ito, C., Moore, C.M., Reddi, A.R. and Williams, L.D. (2020) Human ribosomal G-quadruplexes regulate heme bioavailability. J Biol Chem, 295, 14855–14865.

40. Mestre-Fos, S., Penev, P.I., Suttapitugsakul, S., Hu, M., Ito, C., Petrov, A.S., Wartell, R.M., Wu, R.H. and Williams, L.D. (2019) G-Quadruplexes in Human Ribosomal RNA. Journal of Molecular Biology, 431, 1940–1955.

41. Ivanov, P., O’Day, E., Emara, M.M., Wagner, G., Lieberman, J. and Anderson, P. (2014) G-quadruplex structures contribute to the neuroprotective effects of angiogenin-induced tRNA fragments. P Natl Acad Sci USA, 111, 18201–18206.

42. Lyons, S.M., Gudanis, D., Coyne, S.M., Gdaniec, Z. and Ivanov, P. (2017) Identification of functional tetramolecular RNA G-quadruplexes derived from transfer RNAs. Nature Communications, 8.

43. Kharel, P., Balaratnam, S., Beals, N. and Basu, S. (2020) The role of RNA G-quadruplexes in human diseases and therapeutic strategies. Wiley Interdiscip Rev RNA, 11, e1568.

44. Sanchez de Groot, N., Armaos, A., Grana-Montes, R., Alriquet, M., Calloni, G., Vabulas, R.M. and Tartaglia, G.G. (2019) RNA structure drives interaction with proteins. Nat Commun, 10, 3246.

45. Corley, M., Burns, M.C. and Yeo, G.W. (2020) How RNA-Binding Proteins Interact with RNA: Molecules and Mechanisms. Mol Cell, 78, 9–29.

46. Darnell, J.C., Jensen, K.B., Jin, P., Brown, V., Warren, S.T. and Darnell, R.B. (2001) Fragile X mental retardation protein targets G quartet mRNAs important for neuronal function. Cell, 107, 489–499.

47. Schaeffer, C., Bardoni, B., Mandel, J.L., Ehresmann, B., Ehresmann, C. and Moine, H. (2001) The fragile X mental retardation protein binds specifically to its mRNA via a purine quartet motif. EMBO J, 20, 4803–4813.

48. Kenny, P.J., Kim, M., Skariah, G., Nielsen, J., Lannom, M.C. and Ceman, S. (2020) The FMRP-MOV10 complex: a translational regulatory switch modulated by G-Quadruplexes. Nucleic Acids Res, 48, 862–878.

49. Wu, R., Li, L., Bai, Y., Yu, B., Xie, C., Wu, H., Zhang, Y., Huang, L., Yan, Y., Li, X. et al.. (2020) The long noncoding RNA LUCAT1 promotes colorectal cancer cell proliferation by antagonizing Nucleolin to regulate MYC expression. Cell Death Dis, 11, 908.

50. Sexton, A.N. and Collins, K. (2011) The 5’ guanosine tracts of human telomerase RNA are recognized by the G-quadruplex binding domain of the RNA helicase DHX36 and function to increase RNA accumulation. Mol Cell Biol, 31, 736–743.

51. Booy, E.P., Meier, M., Okun, N., Novakowski, S.K., Xiong, S., Stetefeld, J. and McKenna, S.A. (2012) The RNA helicase RHAU (DHX36) unwinds a G4-quadruplex in human telomerase RNA and promotes the formation of the P1 helix template boundary. Nucleic Acids Res, 40, 4110–4124.

52. Procter, J.B., Carstairs, G.M., Soares, B., Mourao, K., Ofoegbu, T.C., Barton, D., Lui, L., Menard, A., Sherstnev, N., Roldan-Martinez, D. et al.. (2021) Alignment of Biological Sequences with Jalview. Methods Mol Biol, 2231, 203–224.

53. Beaudoin, J.D., Jodoin, R. and Perreault, J.P. (2014) New scoring system to identify RNA G-quadruplex folding. Nucleic Acids Res, 42, 1209–1223.

54. Bedrat, A., Lacroix, L. and Mergny, J.L. (2016) Re-evaluation of G-quadruplex propensity with G4Hunter. Nucleic Acids Res, 44, 1746–1759.

55. Garant, J.M., Perreault, J.P. and Scott, M.S. (2017) Motif independent identification of potential RNA G-quadruplexes by G4RNA screener. Bioinformatics, 33, 3532–3537.

56. Chan, C.Y. and Kwok, C.K. (2020) Specific Binding of a d-RNA G-Quadruplex Structure with an l-RNA Aptamer. Angew Chem Int Ed Engl, 59, 5293–5297.

57. Chu, C. and Chang, H.Y. (2018) ChIRP-MS: RNA-Directed Proteomic Discovery. Methods Mol Biol, 1861, 37–45.

58. Hardin, C.C., Perry, A.G. and White, K. (2000) Thermodynamic and kinetic characterization of the dissociation and assembly of quadruplex nucleic acids. Biopolymers, 56, 147–194.

59. Mergny, J.L., Phan, A.T. and Lacroix, L. (1998) Following G-quartet formation by UV-spectroscopy. FEBS Lett, 435, 74–78.

60. Renaud de la Faverie, A., Guedin, A., Bedrat, A., Yatsunyk, L.A. and Mergny, J.L. (2014) Thioflavin T as a fluorescence light-up probe for G4 formation. Nucleic Acids Res, 42, e65.

61. Creacy, S.D., Routh, E.D., Iwamoto, F., Nagamine, Y., Akman, S.A. and Vaughn, J.P. (2008) G4 resolvase 1 binds both DNA and RNA tetramolecular quadruplex with high affinity and is the major source of tetramolecular quadruplex G4-DNA and G4-RNA resolving activity in HeLa cell lysates. J Biol Chem, 283, 34626–34634.

62. Zhang, Z., Kim, T., Bao, M., Facchinetti, V., Jung, S.Y., Ghaffari, A.A., Qin, J., Cheng, G. and Liu, Y.J. (2011) DDX1, DDX21, and DHX36 helicases form a complex with the adaptor molecule TRIF to sense dsRNA in dendritic cells. Immunity, 34, 866–878.

63. Hutchinson, J.N., Ensminger, A.W., Clemson, C.M., Lynch, C.R., Lawrence, J.B. and Chess, A. (2007) A screen for nuclear transcripts identifies two linked noncoding RNAs associated with SC35 splicing domains. BMC Genomics, 8, 39.

64. West, J.A., Davis, C.P., Sunwoo, H., Simon, M.D., Sadreyev, R.I., Wang, P.I., Tolstorukov, M.Y. and Kingston, R.E. (2014) The long noncoding RNAs NEAT1 and MALAT1 bind active chromatin sites. Mol Cell, 55, 791–802.

65. Saitoh, N., Spahr, C.S., Patterson, S.D., Bubulya, P., Neuwald, A.F. and Spector, D.L. (2004) Proteomic analysis of interchromatin granule clusters. Mol Biol Cell, 15, 3876–3890.

66. Fox, A.H., Lam, Y.W., Leung, A.K., Lyon, C.E., Andersen, J., Mann, M. and Lamond, A.I. (2002) Paraspeckles: a novel nuclear domain. Curr Biol, 12, 13–25.

67. Li, L., Williams, P., Ren, W., Wang, M.Y., Gao, Z., Miao, W., Huang, M., Song, J. and Wang, Y. (2021) YY1 interacts with guanine quadruplexes to regulate DNA looping and gene expression. Nat Chem Biol, 17, 161–168.

68. Herviou, P., Le Bras, M., Dumas, L., Hieblot, C., Gilhodes, J., Cioci, G., Hugnot, J.P., Ameadan, A., Guillonneau, F., Dassi, E. et al.. (2020) hnRNP H/F drive RNA G-quadruplex-mediated translation linked to genomic instability and therapy resistance in glioblastoma. Nature Communications, 11.

69. Chu, C., Zhang, Q.C., da Rocha, S.T., Flynn, R.A., Bharadwaj, M., Calabrese, J.M., Magnuson, T., Heard, E. and Chang, H.Y. (2015) Systematic discovery of Xist RNA binding proteins. Cell, 161, 404–416.

70. Sun, Z.Y., Wang, X.N., Cheng, S.Q., Su, X.X. and Ou, T.M. (2019) Developing Novel G-Quadruplex Ligands: from Interaction with Nucleic Acids to Interfering with Nucleic Acid(-)Protein Interaction. Molecules, 24.

71. Abdelhaleem, M., Maltais, L. and Wain, H. (2003) The human DDX and DHX gene families of putative RNA helicases. Genomics, 81, 618–622.

72. Heddi, B., Cheong, V.V., Martadinata, H. and Phan, A.T. (2015) Insights into G-quadruplex specific recognition by the DEAH-box helicase RHAU: Solution structure of a peptide-quadruplex complex. Proc Natl Acad Sci U S A, 112, 9608–9613.

73. Umar, M.I. and Kwok, C.K. (2020) Specific suppression of D-RNA G-quadruplex-protein interaction with an L-RNA aptamer. Nucleic Acids Res, 48, 10125–10141.

74. Ji, D., Lyu, K., Zhao, H. and Kwok, C.K. (2021) Circular L-RNA aptamer promotes target recognition and controls gene activity. Nucleic Acids Res, 49, 7280–7291.

75. Kung, J.T., Colognori, D. and Lee, J.T. (2013) Long noncoding RNAs: past, present, and future. Genetics, 193, 651–669.

76. Fang, Y. and Fullwood, M.J. (2016) Roles, Functions, and Mechanisms of Long Noncoding RNAs in Cancer. Genomics Proteomics Bioinformatics, 14, 42–54.

77. Statello, L., Guo, C.J., Chen, L.L. and Huarte, M. (2021) Gene regulation by long non-coding RNAs and its biological functions. Nat Rev Mol Cell Biol, 22, 96–118.

78. Somarowthu, S., Legiewicz, M., Chillon, I., Marcia, M., Liu, F. and Pyle, A.M. (2015) HOTAIR forms an intricate and modular secondary structure. Mol Cell, 58, 353–361.

79. Liu, F., Somarowthu, S. and Pyle, A.M. (2017) Visualizing the secondary and tertiary architectural domains of lncRNA RepA. Nat Chem Biol, 13, 282–289.

80. Uroda, T., Anastasakou, E., Rossi, A., Teulon, J.M., Pellequer, J.L., Annibale, P., Pessey, O., Inga, A., Chillon, I. and Marcia, M. (2019) Conserved Pseudoknots in lncRNA MEG3 Are Essential for Stimulation of the p53 Pathway. Mol Cell, 75, 982–995 e989.

81. Qian, X., Zhao, J., Yeung, P.Y., Zhang, Q.C. and Kwok, C.K. (2019) Revealing lncRNA Structures and Interactions by Sequencing-Based Approaches. Trends Biochem Sci, 44, 33–52.

82. Kamura, T., Katsuda, Y., Kitamura, Y. and Ihara, T. (2020) G-quadruplexes in mRNA: A key structure for biological function. Biochem Biophys Res Commun, 526, 261–266.

83. Martadinata, H. and Phan, A.T. (2014) Formation of a stacked dimeric G-quadruplex containing bulges by the 5’-terminal region of human telomerase RNA (hTERC). Biochemistry, 53, 1595–1600.

84. Cusanelli, E. and Chartrand, P. (2015) Telomeric repeat-containing RNA TERRA: a noncoding RNA connecting telomere biology to genome integrity. Front Genet, 6, 143.

85. Ji, Q., Zhang, L., Liu, X., Zhou, L., Wang, W., Han, Z., Sui, H., Tang, Y., Wang, Y., Liu, N. et al.. (2014) Long non-coding RNA MALAT1 promotes tumour growth and metastasis in colorectal cancer through binding to SFPQ and releasing oncogene PTBP2 from SFPQ/PTBP2 complex. Br J Cancer, 111, 736–748.

86. Ji, Q., Cai, G., Liu, X., Zhang, Y., Wang, Y., Zhou, L., Sui, H. and Li, Q. (2019) MALAT1 regulates the transcriptional and translational levels of proto-oncogene RUNX2 in colorectal cancer metastasis. Cell Death Dis, 10, 378.

87. Amodio, N., Raimondi, L., Juli, G., Stamato, M.A., Caracciolo, D., Tagliaferri, P. and Tassone, P. (2018) MALAT1: a druggable long non-coding RNA for targeted anti-cancer approaches. J Hematol Oncol, 11, 63.

88. Bond, C.S. and Fox, A.H. (2009) Paraspeckles: nuclear bodies built on long noncoding RNA. J Cell Biol, 186, 637–644.

89. Minard, A., Morgan, D., Raguseo, F., Di Porzio, A., Liano, D., Jamieson, A.G. and Di Antonio, M. (2020) A short peptide that preferentially binds c-MYC G-quadruplex DNA. Chem Commun (Camb), 56, 8940–8943.

90. Wen, C.J., Gong, J.Y., Zheng, K.W., He, Y.D., Zhang, J.Y., Hao, Y.H. and Tan, Z. (2020) Targeting nucleic acids with a G-triplex-to-G-quadruplex transformation and stabilization using a peptide-PNA G-tract conjugate. Chem Commun (Camb), 56, 6567–6570.

91. He, Y.D., Zheng, K.W., Wen, C.J., Li, X.M., Gong, J.Y., Hao, Y.H., Zhao, Y. and Tan, Z. (2020) Selective Targeting of Guanine-Vacancy-Bearing G-Quadruplexes by G-Quartet Complementation and Stabilization with a Guanine-Peptide Conjugate. J Am Chem Soc, 142, 11394–11403.

92. Dumbovic, G., Braunschweig, U., Langner, H.K., Smallegan, M., Biayna, J., Hass, E.P., Jastrzebska, K., Blencowe, B., Cech, T.R., Caruthers, M.H. et al.. (2021) Nuclear compartmentalization of TERT mRNA and TUG1 lncRNA is driven by intron retention. Nat Commun, 12, 3308.

93. Lewandowski, J.P., Dumbovic, G., Watson, A.R., Hwang, T., Jacobs-Palmer, E., Chang, N., Much, C., Turner, K.M., Kirby, C., Rubinstein, N.D. et al.. (2020) The Tug1 lncRNA locus is essential for male fertility. Genome Biol, 21, 237.

94. Zhang, Q., Yang, K., Li, J., Chen, F., Li, Y. and Lin, Q. (2021) Long Noncoding RNA HCG11 Acts as a Tumor Suppressor in Gastric Cancer by Regulating miR-942-5p/BRMS1 Axis. J Oncol, 2021, 9961189.

95. Xu, Y., Zheng, Y., Liu, H. and Li, T. (2017) Modulation of IGF2BP1 by long non-coding RNA HCG11 suppresses apoptosis of hepatocellular carcinoma cells via MAPK signaling transduction. Int J Oncol, 51, 791–800.

96. Wang, Y.G., Liu, J., Shi, M. and Chen, F.X. (2018) LncRNA DGCR5 represses the development of hepatocellular carcinoma by targeting the miR-346/KLF14 axis. J Cell Physiol, 234, 572–580.

97. Xue, C., Chen, C., Gu, X. and Li, L. (2021) Progress and assessment of lncRNA DGCR5 in malignant phenotype and immune infiltration of human cancers. Am J Cancer Res, 11, 1–13.

98. Zhang, L., Zhang, J., Li, S., Zhang, Y., Liu, Y., Dong, J., Zhao, W., Yu, B., Wang, H. and Liu, J. (2021) Genomic amplification of long noncoding RNA HOTAIRM1 drives anaplastic thyroid cancer progression via repressing miR-144 biogenesis. RNA Biol, 18, 547–562.

99. Wang, H., Li, H., Jiang, Q., Dong, X., Li, S., Cheng, S., Shi, J., Liu, L., Qian, Z. and Dong, J. (2021) HOTAIRM1 Promotes Malignant Progression of Transformed Fibroblasts in Glioma Stem-Like Cells Remodeled Microenvironment via Regulating miR-133b-3p/TGFbeta Axis. Front Oncol, 11, 603128.

