## Supporting information for "Identification and targeting of G-quadruplex structures in *MALAT1* long non-coding RNA"

**Table S1.** Sequences and G4 prediction score of non-coding rG4s found in rG4-seq library.

**Table S2.** Sequences of oligonucleotides, peptide and protein used in this study.

**Table S3.** Sequences of *MALAT1* probes used in this study.

**Figure S1.** rG4 conservation analysis for other 4 MALAT1 rG4s.

**Figure S2.** MST binding of *MALAT1* rG4_04 with DHX36 protein.

**Figure S3.** EMSA binding of *NEAT1* rG4 with NONO protein.

**Figure S4.** EMSA binding of other 4 *MALAT1* rG4s with NONO protein.

**Figure S5.** Filter binding of *MALAT1* rG4_04, *MALAT1* rG4_09 wildtypes and mutants with NONO protein.

**Figure S6.** NONO (53-312) inhibits *MALAT1* rG4_04-DHX36 interaction.

**Figure S7.** MST binding of *MALAT1* rG4_04 with PDS.

**Figure S8.** MST binding of *MALAT1* rG4_04 with L-Apt.4-1c.

**Figure S9.** RHAU53 interferes with *MALAT1* rG4–NONO interactions.

**Table S1.** Sequences and G4 prediction score of non-coding rG4s found in rG4-seq library.

See excel table uploaded separately.

**Table S2**. Sequences of oligonucleotides, peptide and protein used in this study.

| Name | Sequence (5'–3') |
|---|---|
| *MALAT1* rG4_02 [a] | UGGGAUGGUCUUAACAGGGAAGAGAGAGGGUGGGGGA |
| *MALAT1* rG4_04 [a] | AGGGUGGGCUUUUGUUGAUGAGGGAGGGGA |
| *MALAT1* rG4_05 [a] | AGGGAAGGGAGGGGGUGCCUGUGGGG |
| *MALAT1* rG4_06 [a] | AGGGAUGGGAGGAGGGGGUGGGGC |
| *MALAT1* rG4_09 [a] | AGGGGAGGGAAAGGGGGAAAGCGGGC |
| *MALAT1* rG4_10 [a] | AGUGGCUGAGAGGGCUUUUGGGUGGG |
| *NEAT1*_22619 [a] | GGGAGGGAGGGAGGG |
| *MALAT1* rG4_04 MUT [a] | AG**A**GUG**A**GCUUUUGUUGAUGAG**A**GAG**AA**GA |
| Scramble G-rich sequence [a] | GCGAGUGUGUGAGACGUCGC |
| *MALAT1* rG4_04 [b] | AGGGUGGGCUUUUGUUGAUGAGGGAGGGGA |
| *MALAT1* rG4_04 MUT [b] | AG**A**GUG**A**GCUUUUGUUGAUGAG**A**GAG**AA**GA |
| *MALAT1* rG4_09 [b] | AGGGGAGGGAAAGGGGGAAAGCGGGC |
| *MALAT1* rG4_09 MUT [b] | AG**AA**GAG**A**GAAAG**AAA**GAAAGCG**A**GC |
| *MALAT1* forward primer 1 | GCTCTGTGGTGTGGGATTGA |
| *MALAT1* reverse primer 1 | GTGGCAAAATGGCGGACTTT |
| *MALAT1* forward primer 2 | CAGCAGCAGACAGGATTCCA |
| *MALAT1* reverse primer 2 | TCGTTAGCGCTCCTTCCTTC |
| *Xist* forward primer | GGTGCAGGGCTTAAAATGGC |
| *Xist* reverse primer | GTCCACCACCATGCTAACCA |
| *GAPDH* forward primer | GAAAGTAGGGCCCGGCTAC |
| *GAPDH* reverse primer | GTGACCAGGCGCCCAATAC |
| L-Apt.4-1c [c] | GCCCUAAAGGUGGUGGUGGGAGGGC |
| RHAU53 | SMHPGHLKGREIGMWYAKKQGQKNKEAERQERAVVHMDERREEQI VQLLNSVQAK |

| | |
|---|---|
| NONO (53-312) | MHHHHHHEGL TIDLKNFRKP GEKTFTQRSR LFVGNLPPDI TEEEMRKLFE KYGKAGEVFI HKDKGFGFIR LETRTLAEIA KVELDNMPLR GKQLRVRFAC HSASLTVRNL PQYVSNELLE EAFSVFGQVE RAVVIVDDRG RPSGKGIVEF SGKPAARKAL DRCSEGSFLL TTFPRPVTVE PMDQLDDEEG LPEKLVIKNQ QFHKEREQPP RFAQPGSFEY EYAMRWKALI EMEKQQQDQV DRNIKEAREK LEMEMEAARH EHQVMLM |

[a] FAM attached to the 5′ end of all oligos.[b] Biotin attached to the 5′ end of all oligos. [c] All L-RNA bases. The texts in bold indicates G->A mutation.

**Table S3.** Sequences of *MALAT1* probes used in this study.

| Probe No. | Sequence (5'–3') |
|---|---|
| 1 | CTAAATACCACCACCTGGAA |
| 2 | ACACCCAGAAGTGTTTACAC |
| 3 | CAAGGCAAATCGCCATGGAA |
| 4 | CTTGGAAAACGCCTCAATCC |
| 5 | CCTCTTAAAGCACTTCTTGT |
| 6 | GCGAGGCGTATTTATAGACG |
| 7 | GCCCTTCTATTGGTATTAAT |
| 8 | CGTCATGGATTTCAAGGTCT |
| 9 | TCTCCAAATTGTTTCATCCT |
| 10 | ATCTTCTCAAGCTTTACCTT |
| 11 | TACTTCCGTTACGAAAGTCC |
| 12 | CTGGGTCAGCTGTCAATTAA |
| 13 | CTTCACCACCAAATCGTTAG |
| 14 | CCCATATAAATCCCTTTACA |
| 15 | GCCTTTAGGATTCTAGACAG |
| 16 | CTCAACGTGAGAACTGCTCT |
| 17 | CTCTAACCCAGTTTGTCAAT |
| 18 | TGAACCAAAGCTGCACTGTG |
| 19 | ACTGCCAACTAATTGCCAAT |
| 20 | ACTTTCCTTGCCCAAATTAA |
| 21 | ATTGTAGTTAATGTCAGCCC |
| 22 | CTTCAGGATCATTAAGCCAC |
| 23 | CACCCTCTAAGAGACATTCA |
| 24 | CCCAATGGAGGTATGACATA |
| 25 | CATGCAATACTGCAGATGCA |
| 26 | TTTCTCAATCCTGAAATCCC |
| 27 | CATCAAGGCACTGATCACTT |
| 28 | TCCTGATCTGGTCCATTAAA |
| 29 | CTTATTTAGAGGGCCTCTAT |
| 30 | AACATTGCCTACCACTCTAA |
| 31 | CCTGAATGGCTTCATGAAGG |
| 32 | TGCATTTACTTGCCAACAGA |
| 33 | CAACACTCAGCCTTTATCAC |
| 34 | CTGTTGCTTGTTTGGAATGT |
| 35 | CCACTGGTGAATTCAACTGG |
| 36 | AACACAGTTTGCTCACATGC |
| 37 | ATGGTTGTCCTACTTTAAGC |
| 38 | CACTCCAGAAAGAGGGAGTT |
| 39 | GCTTGAGATTTGGGCTTTAT |
| 40 | TTGCAGGCAAATTAATGGCC |

Note: All the probes are 5' biotin-labelled.

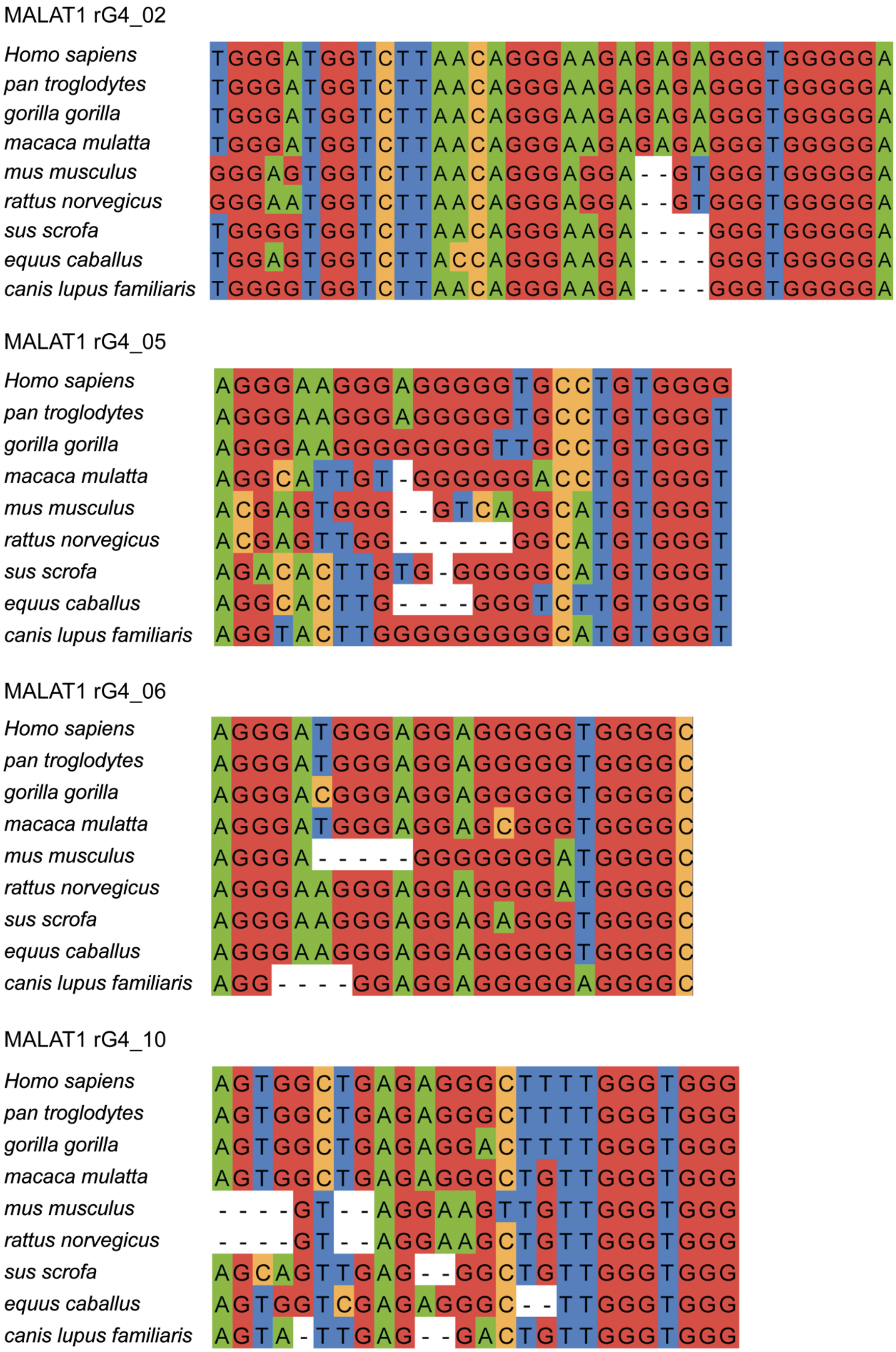


**Figure S1.** rG4 conservation analysis for other 4 MALAT1 rG4s. Comparative sequence analysis of *MALAT1* rG4_02, *MALAT1* rG4_05, *MALAT1* rG4_06 and *MALAT1* rG4_10 in different species.

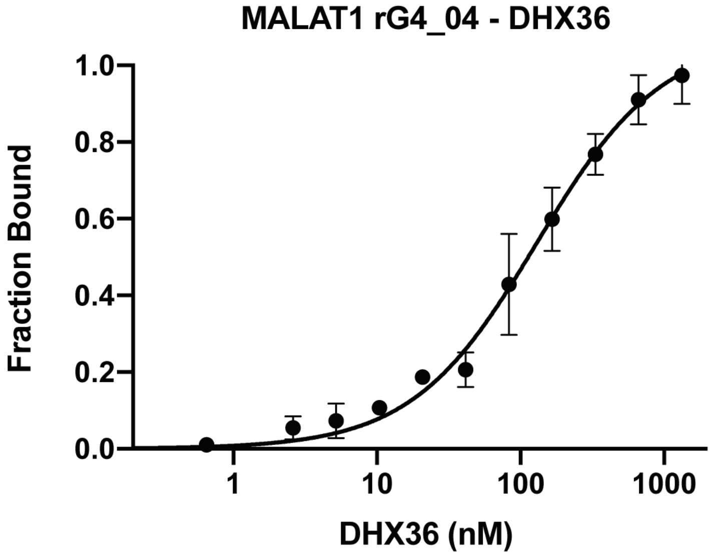


**Figure S2.** MST binding of *MALAT1* rG4_04 with DHX36 protein. EMSA with purified DHX36 (0-1300 nM) and *MALAT1* rG4_04 (fixed at 50 nM). The result suggests the interaction of *MALAT1* rG4_04 with DHX36 protein. The Kd is determined to be 130 ± 16 nM.

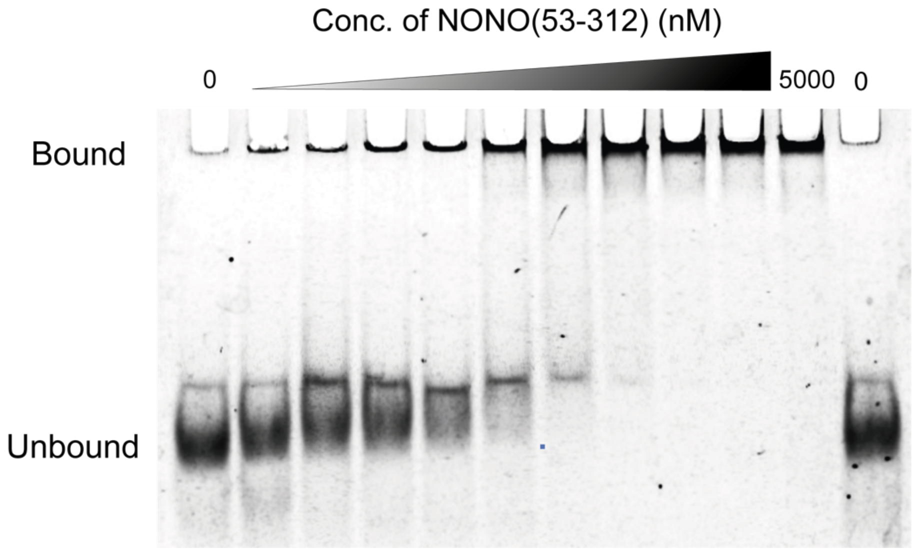


**Figure S3.** EMSA binding of *NEAT1* rG4 with NONO protein. EMSA with recombinant NONO (53-312) (0-5000 nM) and *NEAT1*_22619 (fixed at 5 nM). From the gel, the free *NEAT1* rG4 unbound band gradually decreases with the increasing NONO protein concentration, while the rG4-NONO complex band increases and saturates, suggesting the interaction of *NEAT1* rG4 with NONO protein.

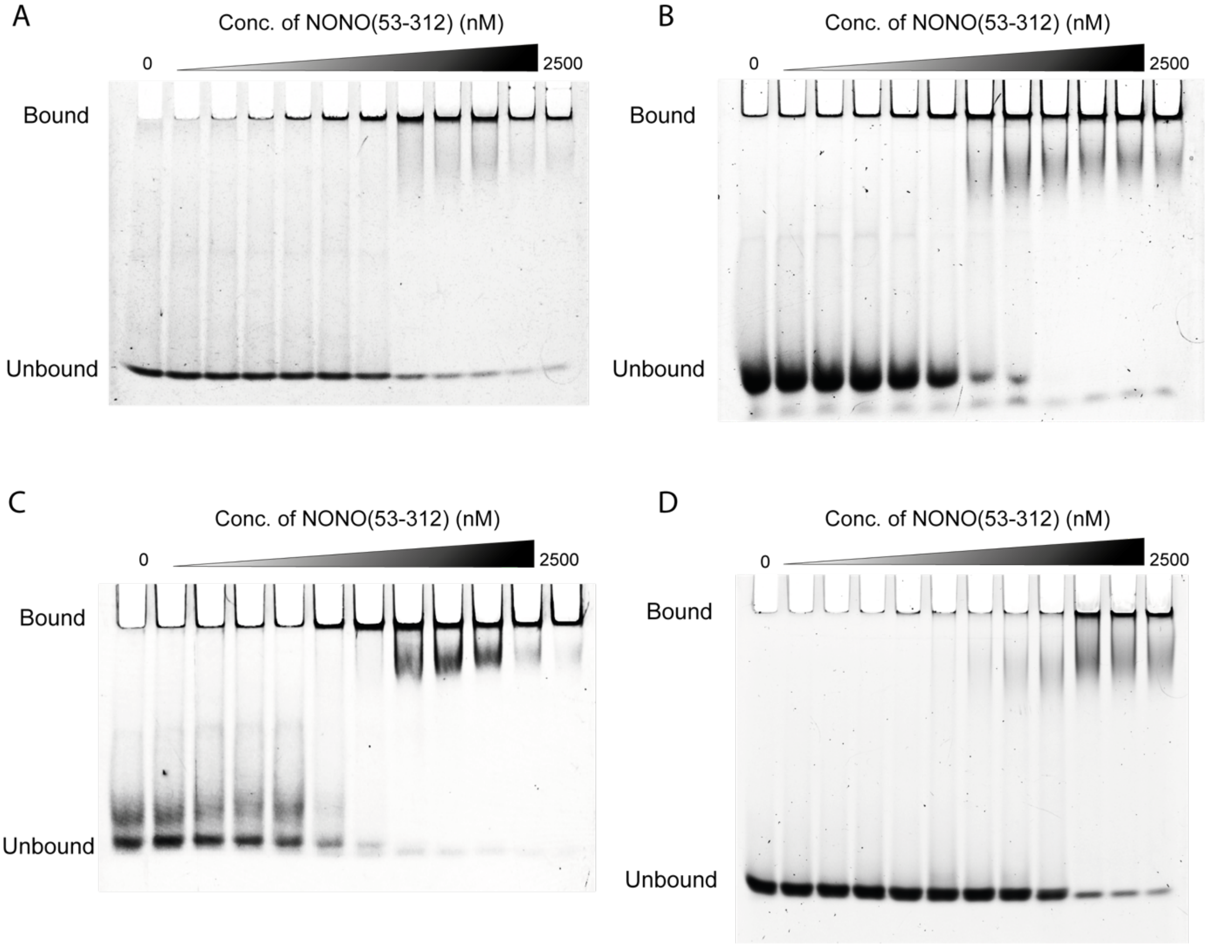


**Figure S4.** EMSA binding of other 4 *MALAT1* rG4 with NONO protein. **(A)** EMSA with recombinant NONO (53-312) (0-2500 nM) and *MALAT1* rG4_02 (fixed at 5 nM). **(B)** EMSA with recombinant NONO (53-312) and *MALAT1* rG4_05 (fixed at 5 nM). **(C)** EMSA with recombinant NONO (53-312) and *MALAT1* rG4_06 (fixed at 5 nM). **(D)** EMSA with recombinant NONO (53-312) and *MALAT1* rG4_10 (fixed at 5 nM). All 4 *MALAT1* rG4s interact with NONO protein, with Kd of 187.7 ± 24.3 nM, 108.2 ± 26.3 nM, 67.9 ± 21.0 nM, 162.0 ± 37.8 nM for *MALAT1* rG4_02, *MALAT1* rG4_05, *MALAT1* rG4_06, and *MALAT1* rG4_10, respectively.

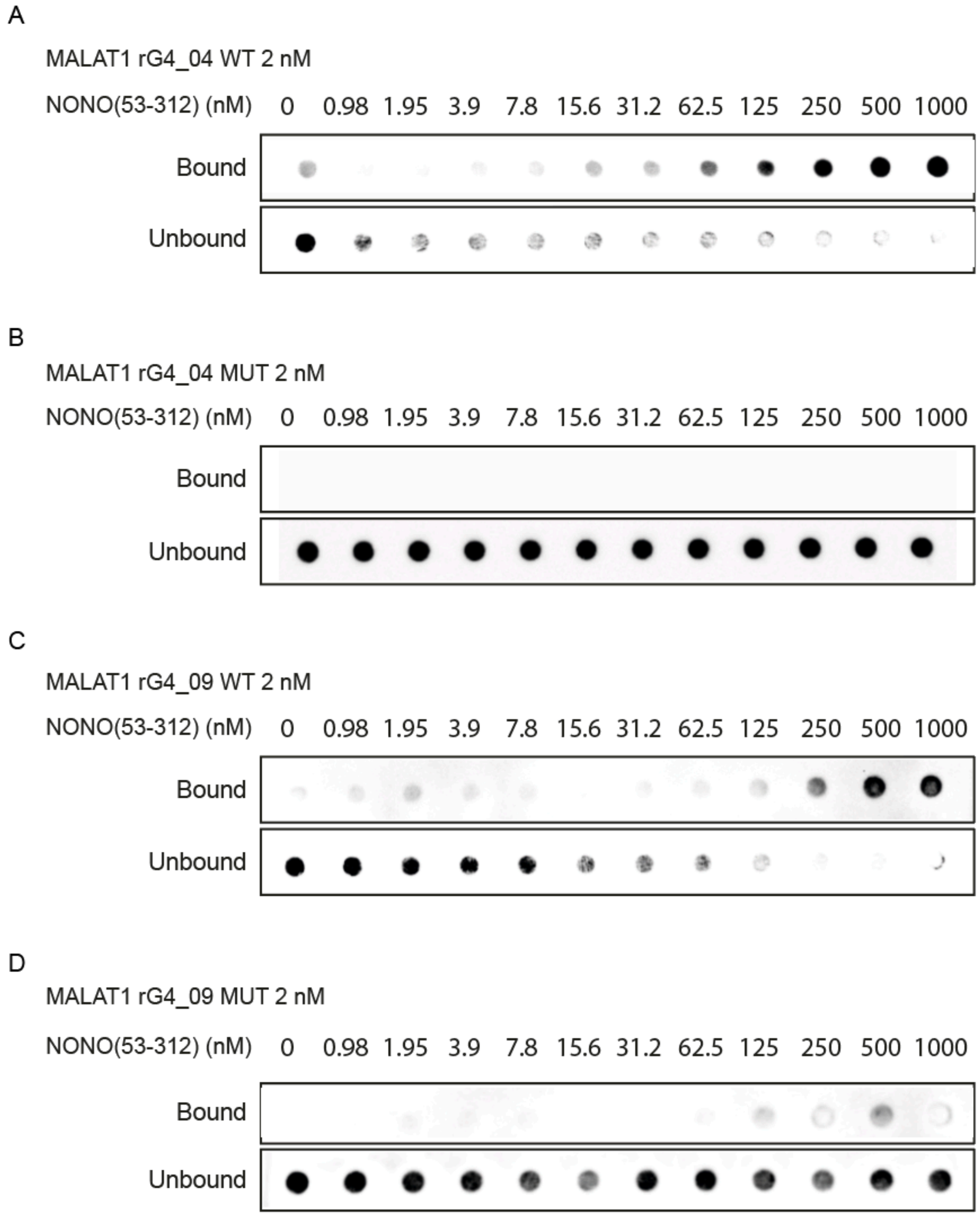


**Figure S5.** Filter binding of *MALAT1* rG4_04, *MALAT1* rG4_09 wildtypes and mutants with NONO protein. **(A)** Filter binding with NONO (53-312) (0-1000 nM) and *MALAT1* rG4_04 wildtype (fixed at 2 nM). **(B)** Filter binding with NONO (53-312) and *MALAT1* rG4_04 mutant. **(C)** Filter binding with NONO (53-312) and *MALAT1* rG4_09 wildtype. **(D)** Filter binding with NONO (53-312) and *MALAT1* rG4_09 mutant.

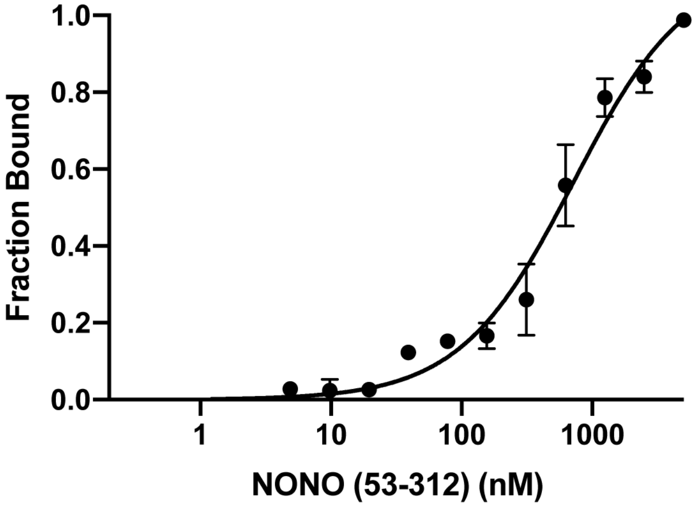


**Figure S6.** NONO (53-312) inhibits *MALAT1* rG4_04-DHX36 interaction. MST with NONO (53-312) (0-5000 nM), DHX36 (fixed at 200 nM) and *MALAT1* rG4_04 (fixed at 50 nM). IC50 was determined to be 718 ± 121 nM. The error bar represents the standard deviation of three independent replicates.

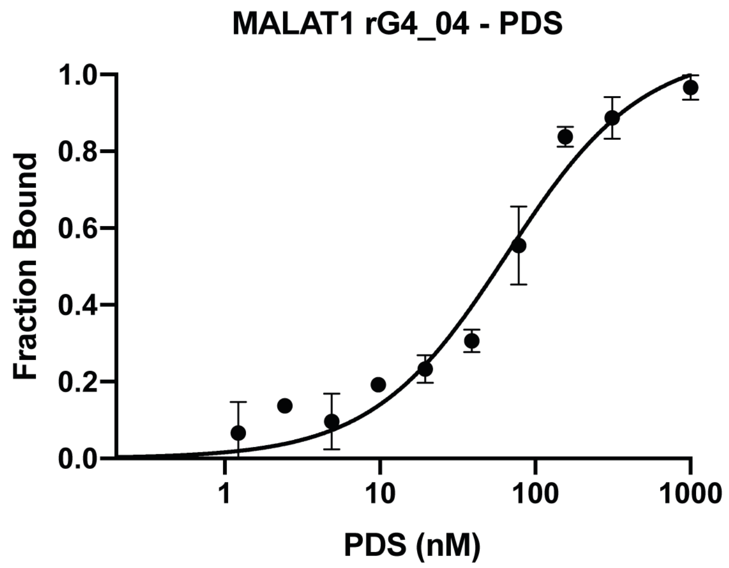


**Figure S7.** MST binding of *MALAT1* rG4_04 with PDS. MST with PDS (0-1000 nM) and *MALAT1* rG4_04 (fixed at 50 nM). The result suggests the interaction of *MALAT1* rG4_04 with PDS. The Kd is determined to be 66 ± 8 nM.

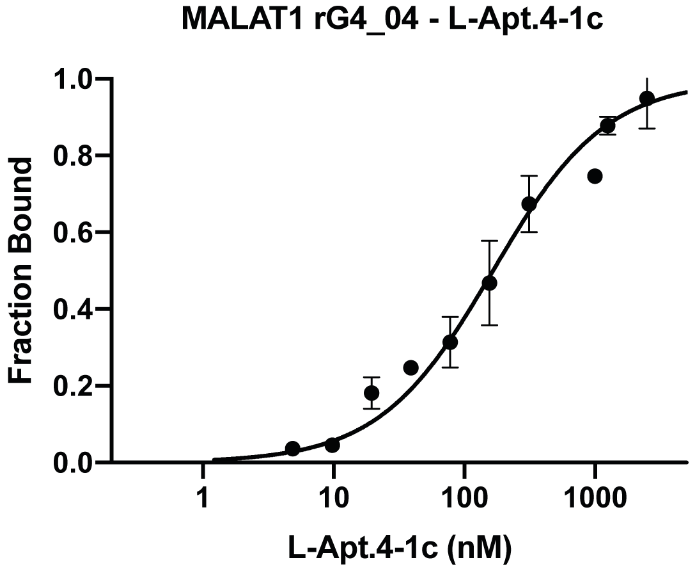


**Figure S8.** MST binding of *MALAT1* rG4_04 with L-Apt.4-1c. MST with L-Apt.4-1c (0-5000 nM) and *MALAT1* rG4_04 (fixed at 50 nM). The result suggests the interaction of *MALAT1* rG4_04 with L-Apt.4-1c. The Kd is determined to be 165 ± 45 nM.

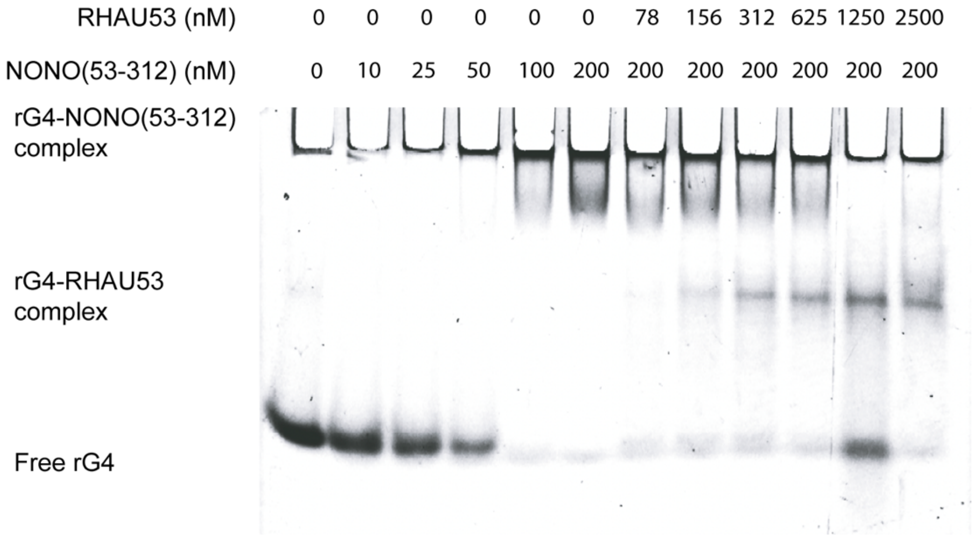


**Figure S9.** RHAU53 interferes with *MALAT1* rG4–NONO interactions. Each lane contained 5 nM FAM MALAT1 rG4_04. In the presence of increasing concentration of RHAU53, the rG4–NONO complex diminishes while the rG4–RHAU53 complex appears.
